# Distributed range adaptation in human parietal encoding of numbers

**DOI:** 10.1101/2025.09.25.675916

**Authors:** Arthur Prat-Carrabin, Gilles de Hollander, Saurabh Bedi, Samuel J. Gershman, Christian C. Ruff

## Abstract

The brain’s representations are encoded in the collective activity of neural populations, whose non-monotonic tuning properties define the population code. Adaptive behavior calls for flexible retuning of this code as stimuli statistics change across contexts. Yet how such adaptation occurs is unknown. Using fMRI during a numerosity-estimation task with variable number ranges, we show that number representations in human parietal cortex dynamically recalibrate to context, enabling context-sensitive behavioral accuracy. The tuning properties of neural populations collectively shift and rescale with the range. This distributed range adaptation achieves efficient coding in real time: neural precision scales with the range and predicts corresponding changes in behavioral precision. Individuals with stronger neural adaptation show larger behavioral adjustments. These findings extend static sensory efficient coding to adaptive representations of abstract magnitudes. Such flexible population tuning may constitute a canonical mechanism of encoding networks that enables the brain to sustain precise, adaptive behavior.

---

Efficient coding prescribes that the brain’s representational resources should be optimally allocated across the stimuli that one may encounter^1–4^. Accordingly, neural tuning curves in various sensory systems are efficiently adapted to the statistics of the sensory stimuli that they encode^5,6^. Yet in natural environments, stimulus statistics typically fluctuate over time and across contexts. For coding to remain efficient, neural representations must therefore be flexible and adapt to these changes. One form of such flexibility is ‘range adaptation’, whereby neurons with monotonic tuning modulate their gains to effectively map the current stimulus range onto their limited response range^7–10^. However, across many species and sensory modalities, the most widespread form of representation relies on populations of neurons with non-monotonic, bell-shaped tuning curves (e.g., orientations^11^, motion direction^12^, spatial frequency^13^, sound frequency^14^, whisker angle^15^, and numerosity^16^, among others^17^). The notion of monotonic range adaptation has no clear analogue in these distributed codes. Hence, although such networks are the predominant form of neural representation, it is largely unknown whether and how they adapt to changes in stimulus statistics.

There is, however, some evidence that distributed neural representations are not static: experimental studies have exhibited shifts of neuronal tuning curves, when a stimulus is the locus of attention or when it is repeatedly presented, and in the encoding of reward intervals^18–24^. Here we propose that this flexibility more generally enables neural populations to adapt to changing stimulus statistics. Specifically, we hypothesize that when the range of possible stimuli changes, the receptive fields of the encoding neural populations shift and scale in a way that efficiently remaps neural representations to cover the new range. We call this mechanism ‘distributed range adaptation’. It involves the concerted adaptation of the entire sensory network to a change in the range of stimuli. This neural mechanism could be a property shared by many sensory circuits, enabling the flexible allocation of processing resources to achieve optimal sensitivity, in line with the principle of efficient coding.

Here we examine our proposal in the context of number representation in humans. Previous research on neural tuning and perception has often focused on the sensory encoding of physical attributes of stimuli (e.g., orientations). Yet many natural decisions are based on abstract quantities that are not directly rooted in physical magnitudes^17,25–28^. In particular, the ability to represent and act upon numerical information is a fundamental skill shared across a wide evolutionary range of species, from primates to squids^29–31^. In humans, this ‘number sense’ is supported by number-sensitive neurons that exhibit bell-shaped receptive fields centered on their preferred numerosities^31–37^. Recent behavioral evidence shows that the perception of numbers by humans exhibits larger errors when the range of possible numbers in a task increases, suggesting a dynamic and efficient adaptation of number representations^38–41^. How this reflects a reallocation of encoding resources at the neural level is unknown.

Encouragingly, numerosity tuning strength in parietal cortex varies with task demands^42^ and individual differences in tuning acuity relate to individual differences in numerical decision-making^43,44^. This suggests that these responses reflect representations supporting numerical perception and decision-making. Adaptive adjustments of parietal numerosity encoding to changes in stimulus statistics — including shifts of preferred numerosities^22,45^ — have moreover been reported^46^, making number representation a strong candidate for probing distributed range adaptation in humans. However, a direct link between these neural adaptations and the principle of efficient coding, as well as their relationship to behavioral changes, have remained missing. Here, we fill this gap by characterizing how the neural mechanism of distributed range adaptation underlies these effects. To this end, we use forward encoding models and behavioral measurements to quantify how tuning curves rescale with stimulus range, how this optimizes representational precision, and how this predicts individual variability in numerical perception.

We investigate our proposal in 39 participants tasked with estimating the number of dots in visual displays while undergoing 3T functional MRI, as we manipulate the range of the distribution of numbers (the prior). Neurocomputational modeling of the fMRI data with numerical population receptive field (nPRF) models^33^ enables us to identify tuning properties of numerosity-selective populations in the intraparietal sulcus (IPS), quantify their changes across priors, and test our hypotheses. We find that the preferred numerosities of the neural populations shift across priors in a way that closely aligns with the quantitative predictions of our proposal. In addition, the widths of the population receptive fields broaden with the width of the prior. Across priors, the encoding populations thus reorganize in a structured fashion to efficiently cover the relevant range, consistent with distributed range adaptation.

Identifying the tuning properties of neural populations also enables us to quantify the representational acuity of the neural population code. With a wider prior, we find that the neural encoding is less precise, consistent with the hypothesis that distributed range adaptation implements efficient coding. The fMRI data, moreover, correlates with individual variations in behavior. Specifically, the individual changes in neural encoding across priors correlate with the individual changes in response precision, supporting the behavioral relevance of the dynamic efficient neural coding evident in the parietal populations. Finally, we also show that within the context of a given prior, the neural data correlate with individual variations in how participants compress larger numbers (Weber’s law): participants whose neural precision decreases faster with large numbers provide less precise estimates for these numbers.

Overall, our results extend the principles of sensory efficient coding to the dynamic adaptation of abstract numerical representations in humans, mediated by distributed range adaptation as a possible canonical mechanism of sensory circuits. Our work also highlights how encoding models for fMRI can be leveraged to probe the neural substrates of cognitive representations, and to establish links, at the individual level, between neural activity and decision-making.

## Behavioral variability and distributed range adaptation

On each trial of a numerosity-estimation task conducted in an MRI scanner, we present participants with a cloud of dots for 600ms and ask them to provide their best estimate of the number of dots (Fig. 1a), enabling us to measure their response variability (Fig. 1b). The number in each trial is randomly sampled from a uniform prior, whose width differs in two experimental conditions. In the ‘Narrow’ condition, the number of dots ranges from 10 to 25, while in the ‘Wide’ condition it ranges from 10 to 40. The prior width in the Wide condition is thus twice as large as in the Narrow condition (Fig. 1c; see Methods).

**Fig. 1:**
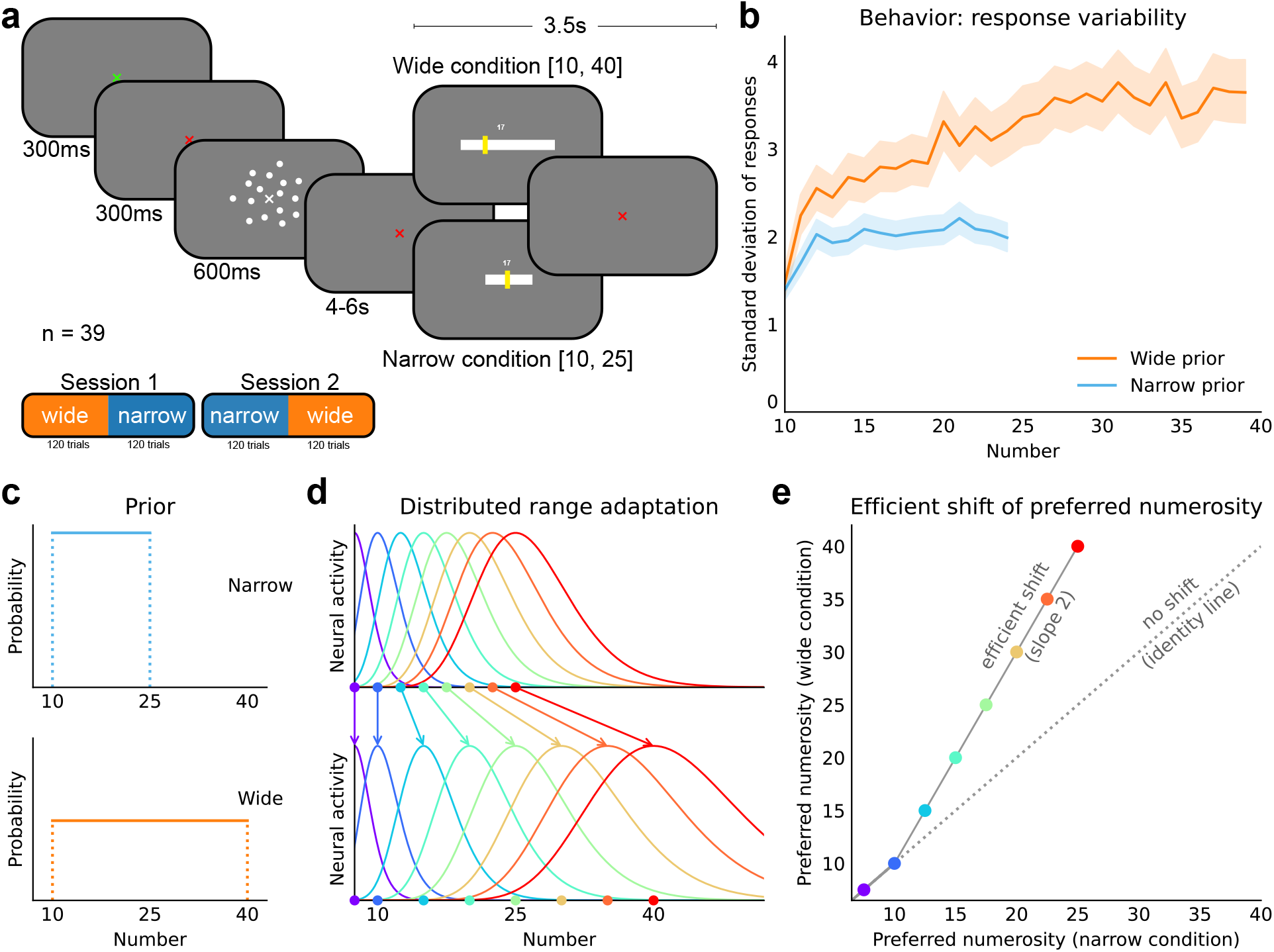
Behavioral task and illustration of distributed range adaptation. **a,** Numerosity-estimation task conducted in the MRI scanner. The participant is asked to estimate the number of dots in a cloud presented for 600ms. This number is randomly sampled from a Narrow or a Wide range (see **c**), and each participant experiences twice each prior across two experimental sessions. **b,** Standard deviation of the participants’ responses as a function of the correct number, in the two conditions. Participants are more variable with the Wide prior. Shaded areas show the 5%-95% credible intervals (see Methods). **c,** Narrow and Wide uniform priors used in the two conditions of the experiment. The Wide prior (10-40) is twice as large as the Narrow prior (10-25). **d,** Schematic illustration of distributed range adaptation: the populations’ receptive fields in the Narrow condition (top panel) are re-allocated in the Wide condition (bottom panel), to efficiently cover the wider range. Hence the receptive fields shift and widen, and over the Narrow range the preferred numerosities in the Wide vs the Narrow condition describe a line of slope 2, the ratio of the prior widths (**e**).

The variability in participants’ estimates is greater in the Wide condition than in the Narrow condition (Fig. 1b), replicating previous results^38^ (the participants’ average estimates are reported in Supplementary Information). This increased variability suggests a more imprecise representation of the numbers with the Wide prior. In turn, this points to an adaptive and dynamic implementation of efficient coding in which the allocation of representational resources optimally adjusts to the current range of numbers, resulting in lower precision for each number under the Wide prior.

We hypothesize that the observed changes in behavioral variability reflect changes in the neural code that dynamically implement efficient coding. Numerosity-sensitive neural populations have bell-shaped receptive fields characterized by their preferred numerosities and their widths^33,47^. Collectively, these tuning properties define the precision of the encoding network for the different stimuli. Efficient-coding models prescribe that this precision should match the prior, i.e., more frequent stimuli should be encoded with greater precision^3,4^. Yet these models typically do not specify how the encoding should transition across priors: in other words, it is unclear what the tuning properties under one prior should become under another prior.

We thus formulate a hypothesis regarding the changes in tuning. Several efficient-coding models predict that, given a prior, the density of the preferred stimuli of encoding neurons should be proportional to this prior^3,48,49^. Integrating this density implies that the cumulative fraction of neurons tuned to stimuli below a stimulus *x*, *D*(*x*), is the cumulative distribution function (CDF) of the prior, *F* (*x*). In other words, a neuron’s relative position (rank) in the ordered population corresponds to a quantile of the prior. For numerosity and many other stimuli, moreover, neural populations are topographically organized, i.e., the ordering of neurons according to their preferred stimuli correlates with their spatial arrangement^33^. We thus assume that when the prior changes, the neurons collectively reorganize their response profiles so as to maintain their relative positions in the distribution. Thus, across different priors, neurons should remain tuned to the same quantiles. Formally, denoting by *µ*_w_ and *µ*_n_ the preferred numerosities of a neuron in the Wide and Narrow conditions, our hypothesis is that

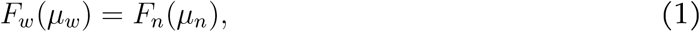

where *F*_w_ and *F*_n_ are the CDFs of the Wide and Narrow priors, respectively.

As uniform priors have linear CDFs, this implies, for preferred numerosities in the Narrow range (*µ*_n_ ∈ [10, 25]), a linear relationship:

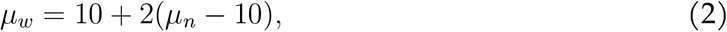

where the slope 2 is the ratio of the prior widths. Under this hypothesis, two preferred numerosities in the Narrow range and separated by *dµ*_n_, in the Narrow condition, are predicted to be separated by twice this distance, *dµ*_w_ = 2 *dµ*_n_, in the Wide condition. Informally, the preferred numerosities thus “spread out” to cover the wider prior, a collective adjustment that we call distributed range adaptation (Fig. 1d). Consistent with efficient coding, less resources are thus dedicated to each number in the Wide condition, resulting in lower precision. Note that distributed range adaptation is not in itself a prediction of efficient coding; rather, it is a proposed mechanism to attain efficient coding flexibly, in sensory networks.

Numbers below 10 have vanishing probabilities under both priors (∀*µ*_n_*, µ*_w_ ≤ 10, *F*_w_(*µ*_w_) = *F*_n_(*µ*_n_) = 0), thus our hypothesis (Eq. 1) does not uniquely determine the relation across conditions between the preferred numerosities in this range. Here we simply conjecture that the preferred numerosities remain stable, i.e., *µ*_n_ = *µ*_w_. We note that this is also the prediction we would make under our hypothesis, if we additionally assumed that participants held subjective priors that were mixtures of the correct prior in each condition and of a constant distribution that allocated non-vanishing probabilities to numbers below 10, such as a long-term prior. However, in our analyses, we also examine the alternative possibility that preferred numerosities scale by the same ratio everywhere, i.e., *µ*_w_ = 2*µ*_n_ (see Supplementary Information).

To illustrate, if a population’s preferred numerosity is 15 in the Narrow condition, our hypothesis (Eq. 1) is thus that it will shift to 20 in the Wide condition. We call such changes in preferred numerosities ‘efficient shifts’ (Fig. 1e). This ‘efficient-shift’ relationship is a quantitative prediction that puts a strong constraint on the receptive fields we should observe across priors.

Finally, regarding the widths of the receptive fields, some efficient-coding models posit a ‘tiling’ property that postulates an inverse relationship between receptive-field density and width^3,49^. Thus one might expect the widths to double in size in our experiment, following the doubling of the prior range. But if both the density and the widths scale with a factor 2 between the Narrow and Wide conditions (all else being equal), then the imprecision of the encoding, as measured for instance by the standard deviation of estimates, should scale by the same factor. Previous studies have shown, however, that this is not the case; instead, the imprecision decreases sublinearly with the width of the prior, and thus participants are *relatively* more precise with wider priors^38,39^. For this reason, we do not expect the widths to double in size; if they do expand, we conjecture that it should be by a factor lower than 2.

## Numerical population receptive fields

We now turn to the fMRI data to assess our hypotheses. We fit models of numerical population receptive fields (nPRFs) to the single-trial BOLD responses per voxel at the time of the presentation of the stimulus, specifically in the intraparietal sulcus (IPS; Fig. 2). The region of interest (ROI) we focus on is the right numerical parietal cortex (rNPC) identified as having the most pronounced number fields in several previous studies^33,44,47,50,51^ (Fig. 2a; see Methods). Each nPRF model specifies a voxel’s average activity as a unimodal function of the encoded number, parameterized by its preferred numerosity, its width, its baseline activity, and its amplitude (specifically, in line with previous results, we choose Gaussian functions in logarithmic space; see Methods). We fit different nPRF models corresponding to different hypotheses and we report the proportions of voxels in the ROI with positive cross-validated variance explained (cv*R*^2^ *>* 0; nPRF parameters are only reported for such ‘signal’ voxels). We use cv*R*^2^ as a voxelwise model comparison technique because it balances model flexibility and generalizability with minimal assumptions^52^. By evaluating performance on held-out data, cv*R*^2^ automatically penalizes overfitting, ensuring that improvements in fit from additional parameters generalize to unseen data. Conversely, improvements in fit with fewer parameters suggests that the implied constraints are verified in data. See Supplementary Information for examples of model fits.

**Fig. 2:**
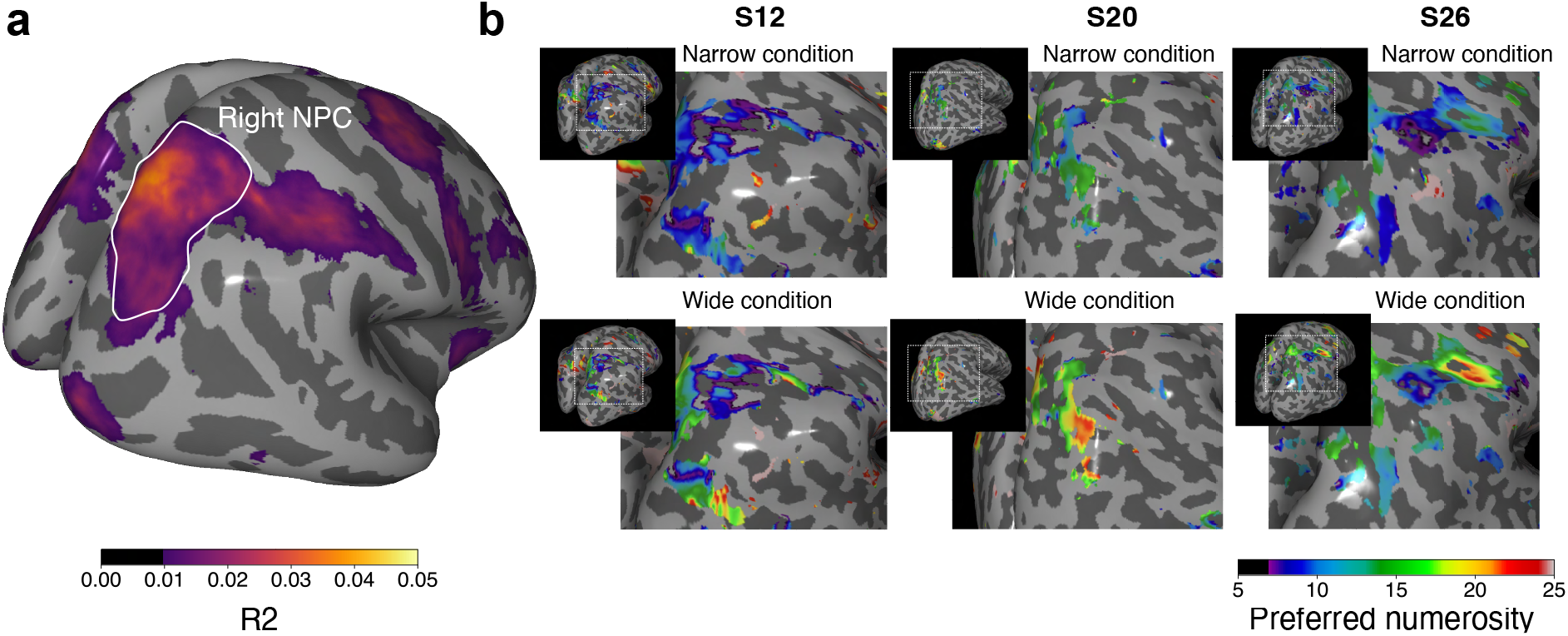
Maps of numerical population receptive fields show shifts of voxels’ preferred numerosities. **a,** Explained variance (*R*^2^) of the nPRF ‘free-shift’ model across 39 participants in *fsaverage*-space. The outlined area indicates the right NPC mask used in all analyses. **b,** Individual maps of preferred numerosities in the two conditions for three representative participants. From the Narrow to the Wide condition, preferred numerosities generally shift upwards. In the ‘free-shift’ model, there are no constraints on the relationship between the two conditions, and each preferred-numerosity parameter of each voxel in each condition is fitted separately. An interactive 3D cortical surface viewer of individual model fits is available online: https://ruffgroup.github.io/neural_priors_viewers/.

## Efficient shifts of the receptive fields

First, we look at the voxels’ preferred numerosities. Fixing all the other parameters, we let the preferred numerosities of each voxel in the Narrow and Wide conditions be free parameters, *µ*_n_ and *µ*_w_ (‘free-shift’ model). We start with this unconstrained model to assess whether the data support our hypothesis in general (below, we test our hypothesis with a more constrained model). The preferred numerosities in the two conditions are strongly correlated (participants pooled: Spearman’s *ρ* = 0.77, *P <* 10*^−^*^320^*, N* = 8131; across participants: average *ρ* = 0.66, interquartile range (IQR): 0.57-0.80). Thus, the neural populations tuned to larger numbers in the Narrow condition are also tuned to larger numbers in the Wide condition. This supports our hypothesis that preferred numerosities maintain their relative ordering across priors. Also as hypothesized, the maps of preferred numerosities in each condition suggest that for a given voxel, the preferred numerosity in the Wide condition is larger than that in the Narrow condition (as shown in Figure 2b for three representative participants). Indeed, the relation between a voxel’s preferred numerosities across the two conditions is not well described by the identity function, except for small numerosities; most voxels with preferred numerosities above 10 in the Narrow condition shift to higher preferred numerosities in the Wide condition (i.e., *µ*_w_ *> µ*_n_; across-participant t-test that the mean shift *µ*_w_ − *µ*_n_ is zero, for *µ*_n_ ≥ 10: *t*(38) = 5.13, *P* = 9 × 10*^−^*^6^; Fig. 3a,b). This increase is observed in most participants: for 37 out of 39 participants, the across-voxels median preferred numerosity is larger in the Wide condition than in the

**Fig. 3:**
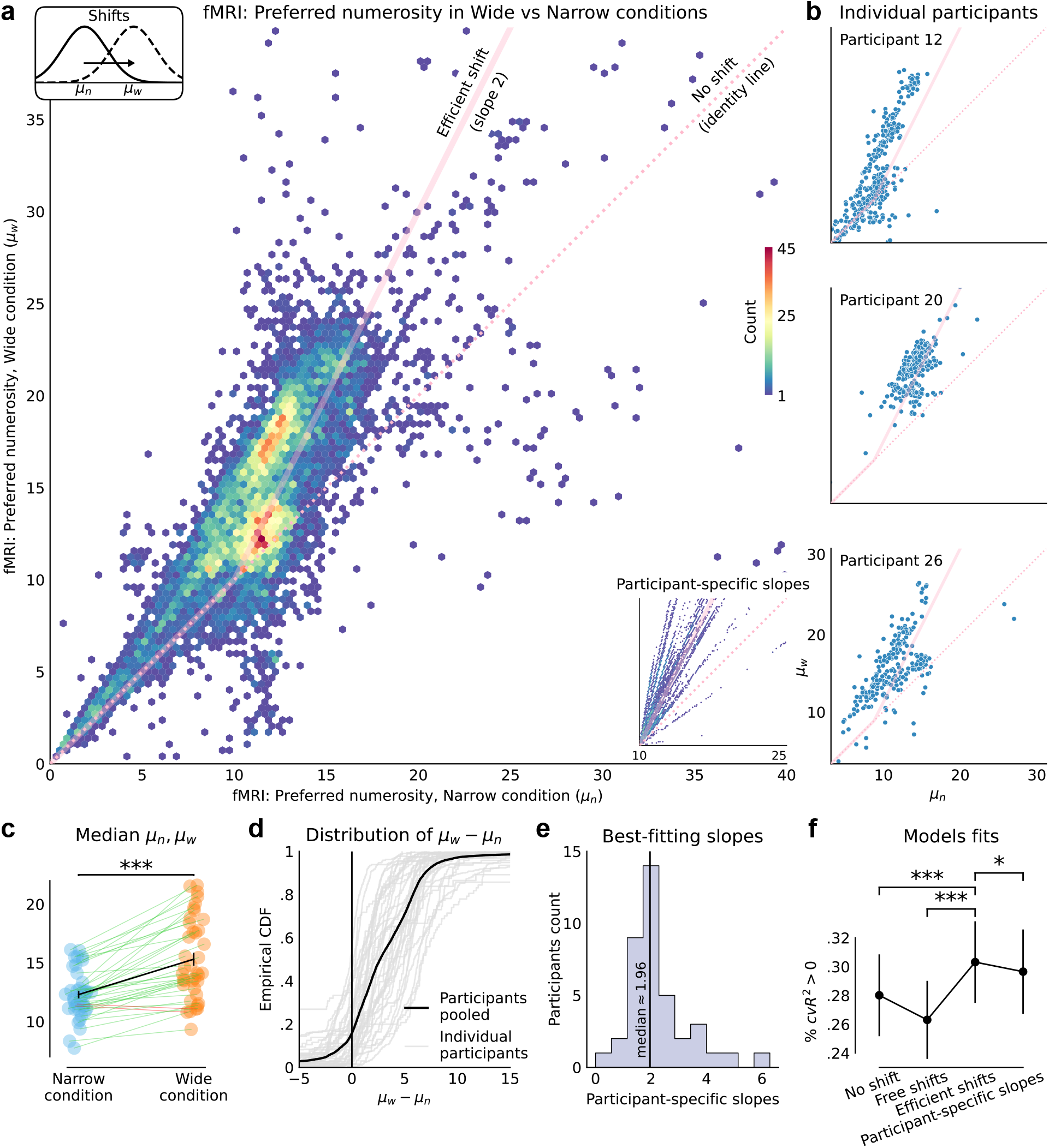
Efficient shifts of neural receptive fields across priors. **a,** Distribution of the voxels’ pairs of best-fitting preferred-numerosity parameters (*µ*_n_*, µ*_w_) in the Wide vs. Narrow conditions (participants pooled, ‘free-shift’ fMRI model). The distribution better aligns with the efficient-shift prediction of distributed range adaptation (transparent pink line) than with the identity function (dotted pink line). 1.94% of outlier voxels not shown (*µ*_n_ or *µ*_w_ *>* 40). *Inset:* Same distribution, with the ‘participant-specific slopes’ model. **b,** Preferred numerosities in the Wide vs. Narrow conditions for three representative participants (‘free-shift’ model). Pink lines as in **a**. **c,** Median preferred numerosities for each participant in the Narrow (left) and Wide (right) conditions. A green (resp., red) line indicates that the participant’s median preferred numerosity in the Wide condition is greater (resp., lower) than in the Narrow condition. Black line shows across-participants averages (error bars: 1 standard error of the mean). **d,** Empirical cumulative distribution of the difference between each voxel’s preferred numerosities in the Wide and Narrow condition, *µ*_w_ − *µ*_n_, for each participant (gray lines) and for the participants pooled (black line). For a majority of voxels this difference is positive, i.e., most voxels’ receptive fields shift towards larger values in the Wide condition. **e,** Distribution of the slope parameter across participants (‘participant-specific slopes’ model). The slope parameter is distributed around 2, the value predicted by distributed range adaptation. **f,** Across-participants average of the proportion of voxels with positive cross-validated variance explained (cv*R*^2^ *>* 0) for (from left to right) the model with no shifts in the voxels’ preferred numerosities, the model with free unrestricted shifts (shown in **a**), the ‘efficient-shift’ model with a slope of 2 (predicted by distributed range adaptation), and the model with participant-specific slopes (inset of **a**). The ‘efficient-shift’ model outperforms the other three. Error bars show ±1 standard error of the mean. ***: *P <* 0.001, **: *P <* 0.01, *: *P <* 0.05.

Narrow condition (Fig. 3c; across-participants paired t-test of equality of the medians in the two conditions: *t*(38) = 8.81, *P* = 1 × 10*^−^*^10^). Overall, the preferred numerosities of 84% of voxels increase in the Wide condition as compared to the Narrow condition (Fig. 3d).

The voxels’ locations across conditions are remarkably well described by the mechanism of distributed range adaptation (compare Fig. 3a,b to Fig. 1e). The preferred numerosities in the Wide vs. Narrow conditions are close to a line of slope 2, as hypothesized, when the preferred numerosity in the Narrow condition is above 10 (we test for this specific relationship with a constrained model, below). For example, voxels whose preferred numerosities are close to 15 in the Narrow condition indeed exhibit preferred numerosities on average close to 20 in the Wide condition (e.g., *µ*_w_ for voxels with *µ*_n_ ∈ [14.5, 15.5] is on average 20.01; standard error of the mean (sem): 0.19). Moreover, we find that the identity function indeed characterizes reasonably well the data when preferred numerosities are below 10 (Fig. 3a,b). (We note however that neither conditions contain numerosities below 10, and thus these estimates should be interpreted cautiously. Further analyses, including parameter recovery, suggest that the encoding is indeed stable across conditions, for these voxels, although we cannot estimate their preferred numerosities as precisely as those above 10; see Supplementary Information). As for larger numbers, only 2.4% of voxels have preferred numerosities greater than 25 in the Narrow condition; due to this scarcity of data, we do not investigate specific hypotheses or constraints that focus on these voxels. To test our hypothesis of efficient shifts, we fit a new model which explicitly enforces the hypothesized relationship between a voxel’s preferred numerosities in the two conditions (i.e., *µ*_w_ = 10 +2(*µ*_n_ − 10) if *µ*_n_ ≥ 10, otherwise *µ*_w_ = *µ*_n_). We emphasize that this is a strong constraint that considerably reduces the number of nPRFs parameters. This ‘Efficient shifts’ model yields, however, a better cross-validated fit than the unconstrained, ‘Free shifts’ model considered thus far (with a significant increase of the proportion of voxels with cv*R*^2^ *>* 0; across-participants paired t-test of equality of the proportions: *t*(38) = 8.30, *P* = 5 × 10*^−^*^10^). It also fits significantly better than a ‘No shift’ model that enforces the identity constraint (*µ*_n_ = *µ*_w_; *t*(38) = 3.73, *P* = 6 × 10*^−^*^4^). Hence from one prior to the other, the preferred numerosities of the encoding neural populations shift in a way that is *quantitatively* consistent with distributed range adaptation. (We also fit alternative models in which *µ*_w_ is proportional to *µ*_n_; they do not yield better fits; see Supplementary Information.)

We also consider the possibility that different participants may adjust differently to the distributions. Specifically, we fit a ‘Participant-specific shifts’ model, similar to the ‘Efficient shifts‘ model, except that the slope is not fixed at 2 but is instead a free parameter for each participant, which we denote by *r*_µ_ (i.e., *µ*_w_ = 10 + *r*_µ_(*µ*_n_ − 10) if *µ*_n_ ≥ 10, else *µ*_w_ = *µ*_n_; see lower-right inset in Fig. 3a). We find that this slope parameter is distributed (across participants) around 2 (median: 1.96, mean: 2.25, sem: 0.19; Fig. 3e), and the mean best-fitting value is significantly different from 1 (t-test *t*(38) = 6.73, *P* = 5.7 × 10*^−^*^8^), but not from 2 (*t*(38) = 1.36, *P* = 0.18). (To test the robustness of our parameter estimation we also examine ‘censored’ fits, restricted to the numbers that occur in both ranges; this analysis supports our conclusions; see Supplementary Information). Moreover, despite this model’s flexibility, it fits significantly worse than the more constrained ‘Efficient shifts‘ model whose slope is fixed at 2 (Fig. 3f; paired t-test *t*(38) = 2.20, *P* = 3.4 × 10*^−^*^2^). In other words, assuming for all the participants a slope of 2, i.e., the value implied by distributed range adaptation, yields a better and more parsimonious account of the data.

## Wider receptive fields with the wider prior

We now turn to the widths of the receptive fields. We start from the ‘efficient shifts’ model and let the widths of each voxel’s receptive field in the two conditions be free parameters, *σ*_n_ and *σ*_w_ (Fig. 4a,b). The two widths are significantly correlated (participants pooled: Spearman’s *ρ* = 0.90, *P <* 10*^−^*^320^, *N* = 8827; across participant: average *ρ* = 0.85, IQR: 0.84-0.94). For a majority of participants (32 out of 39), the median width (across voxels) in the Wide condition is larger than in the Narrow condition (paired t-test of equality: *t*(38) = 6.27, *P* = 2 × 10*^−^*^7^; Fig. 4c). Overall, the widths of 77% of voxels increase in the Wide condition as compared to the Narrow condition (Fig. 4d).

**Fig. 4:**
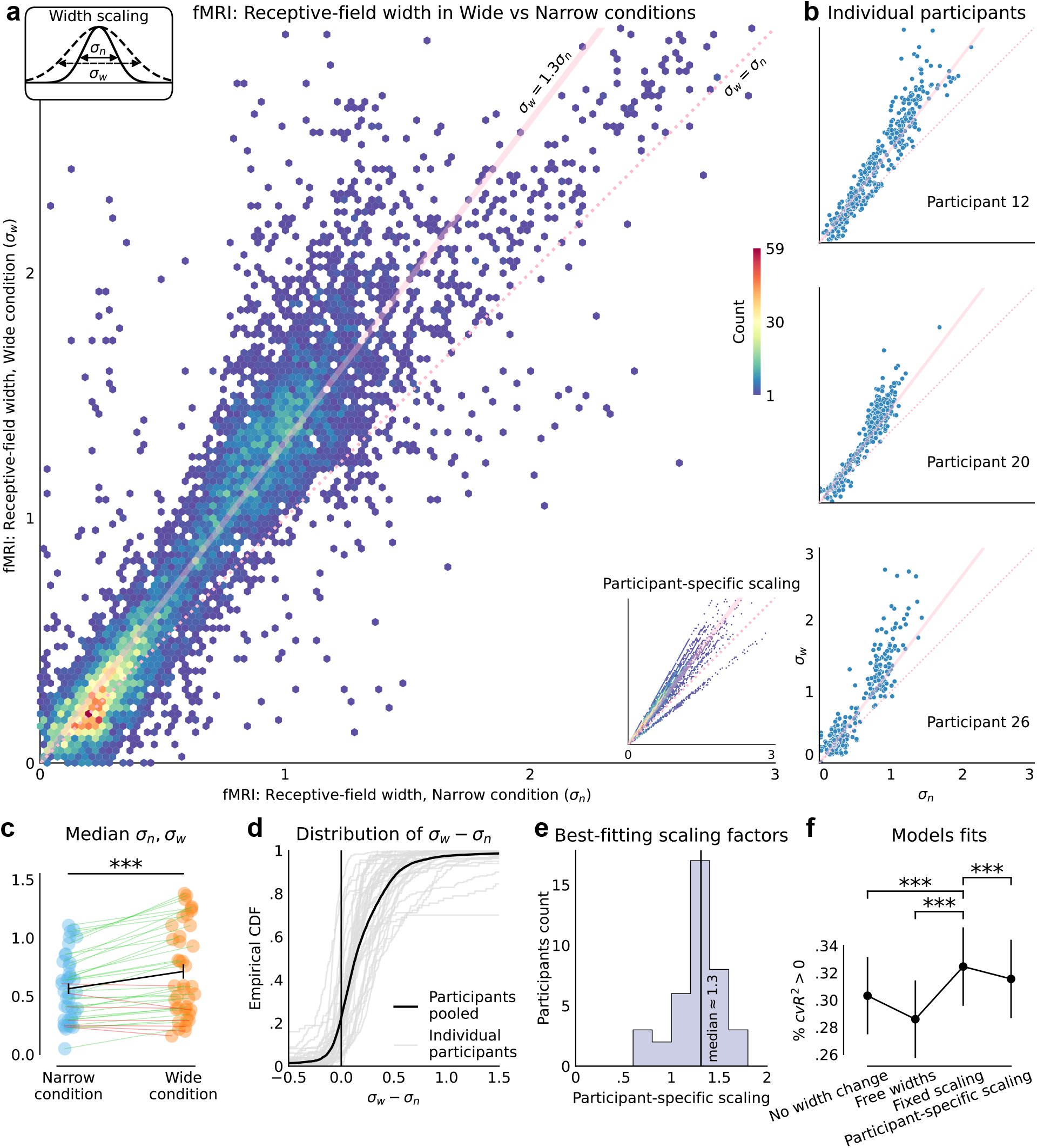
The receptive fields broaden under the wider range. **a,** Distribution of the voxels’ pairs of best-fitting widths parameters (*σ*_n_*, σ*_w_) in the Wide vs. Narrow conditions (participants pooled, ‘free-widths’ model). The widths in the Wide condition are approximately proportional to those in the Wide condition, with a ratio greater than 1, indicating a widening of the receptive fields (transparent pink line: ratio 1.3, dotted pink line: identity function). 2.44% of outlier voxels not shown because one width is outside the figure range. *Inset:* Same distribution, with the ‘participant-specific scaling’ model. **b,** Widths in the Wide vs. Narrow conditions for three representative participants (‘free-widths’ model). Pink lines as in **a**. **c,** Median widths for each participant in the Narrow (left) and Wide (right) conditions. A green (red) line indicates that the participant’s median width in the Wide condition is greater (lower) than in the Narrow condition. Black line shows across-participants averages (error bars: ±1 standard error of the mean). **d,** Empirical cumulative distribution of the difference between each voxel’s widths in the Wide and Narrow condition, *σ*_w_ − *σ*_n_, for each participant (gray lines) and for the participants pooled (black line). For a majority of voxels this difference is positive, i.e., most voxels’ receptive fields widen in the Wide condition. **e,** Distribution of the width-scaling parameter across participants (‘participant-specific scaling’ model). **f,** Across-participants average of the proportion of voxels with positive cross-validated variance explained (cv*R*^2^ *>* 0) for (from left to right) the model with no change in the voxels’ widths parameters, the model with free width parameters (shown in **a**), the ‘fixed width-scaling’ model, and the model with participant-specific scalings (inset of **a**), all with the efficient-shift relation on the preferred numerosities. The ‘fixed width-scaling’ model outperforms the other three. Error bars show ±1 standard error of the mean. ***: *P <* 0.001.

Closer examination of the distribution of the two parameters suggests that the width in the Wide condition is proportional to the width in the Narrow condition, with a scaling factor greater than one (Fig. 4a,b). We thus fit a model in which for each participant a scaling relationship is enforced between the widths of all the voxels in the two conditions, i.e., *σ*_w_ = *r*_σ_*σ*_n_, where *r*_σ_ is a participant-specific scaling parameter. We find that this scaling parameter is significantly greater than 1 (t-test *t*(38) = 7.23, *P* = 1.2 × 10*^−^*^8^), and, as conjectured, significantly lower than 2 (*t*(38) = 17.9, *P* = 4 × 10*^−^*^20^). Its across-participant average is 1.3 (also its median), and its standard deviation is 0.25 (Fig. 4e; we reach the same conclusions when examining ‘censored’ fits restricted to the numbers in the Narrow range; see Supplementary Information).

This relatively low dispersion suggests that the same underlying mechanism operates across participants, prompting us to examine an ‘Efficient shifts, fixed width-scaling’ model in which all the participants have the same scaling factor, *r*_σ_ = 1.3 (which maximizes the model’s likelihood). This yields a significantly better fit than with participant-specific scaling factors (*t*(38) = 5.08, *P* = 10*^−^*^5^), with free width parameters for all the voxels across the two conditions (*t*(38) = 10.5, *P* = 9 × 10*^−^*^13^), or with equal widths across conditions (*σ*_n_ = *σ*_w_; *t*(38) = 4.9, *P* = 1.6 × 10^-5^ ; Fig. 4f). Thus, a better and more parsimonious account of the data is obtained by assuming that from the Narrow to the Wide condition, the widths of the receptive fields scale up by the same factor for all the participants.

This is our best-fitting model of the fMRI data. Consistent with our hypothesis of distributed range adaptation, it is a highly constrained model, that enforces a linear relationship with a specific slope, 2, between the preferred numerosities across conditions, i.e., *µ*_w_ = 10 + 2(*µ*_n_ − 10), for *µ*_n_ ≥ 10, and a linear scaling of the receptive-field widths across conditions, i.e., *σ*_w_ = *r*_σ_*σ*_n_, with *r*_σ_ *>* 1. When examining the amplitudes, we find no evidence that they change across conditions (see Supplementary Information).

## Less precise neural coding with the wider prior

We show substantial changes in the tuning properties of numerosity-sensitive neural populations in the right parietal cortex, across the two conditions (Figs. 2,3,4). Consistent with our hypothesis, receptive fields in the Wide condition are more spread out and broader, a pattern that we argue should reduce encoding precision, in a dynamic implementation of efficient coding. We now test this more directly, by estimating the imprecision of the neural code in the two conditions.

We derive a measure of the encoding precision from the parameters of our nPRF model, combined with a noise component that we estimate by fitting a multivariate distribution to the residuals of the model^53^ (see Methods). We estimate the encoding Fisher information^54^, a statistical measure of precision, for each participant and condition. We find that the inverse of its square-root, 1/ √ *I* (a lower bound on the standard deviation of estimates), is significantly larger in the Wide condition than in the Narrow condition (paired t-test: *t*(38) = 6.53, *P* = 1 × 10*^−^*^7^; Fig. 5a, left panel), indicating reduced precision of neural encoding in the Wide condition. We note, however, that the Fisher information measures sensitivity to infinitesimal changes in the stimulus and, as an approximation to the variance, is only relevant in the small-noise regime of unbiased estimators of continuous quantities. However, in our task the stimulus is discrete, the optimal decoder is Bayesian and therefore biased, and noise levels are not in a small-noise regime. Thus, to better quantify the precision of the neural encoding, we directly estimate it with a simulation-based approach, instead of relying on a theoretical approximation.

**Fig. 5:**
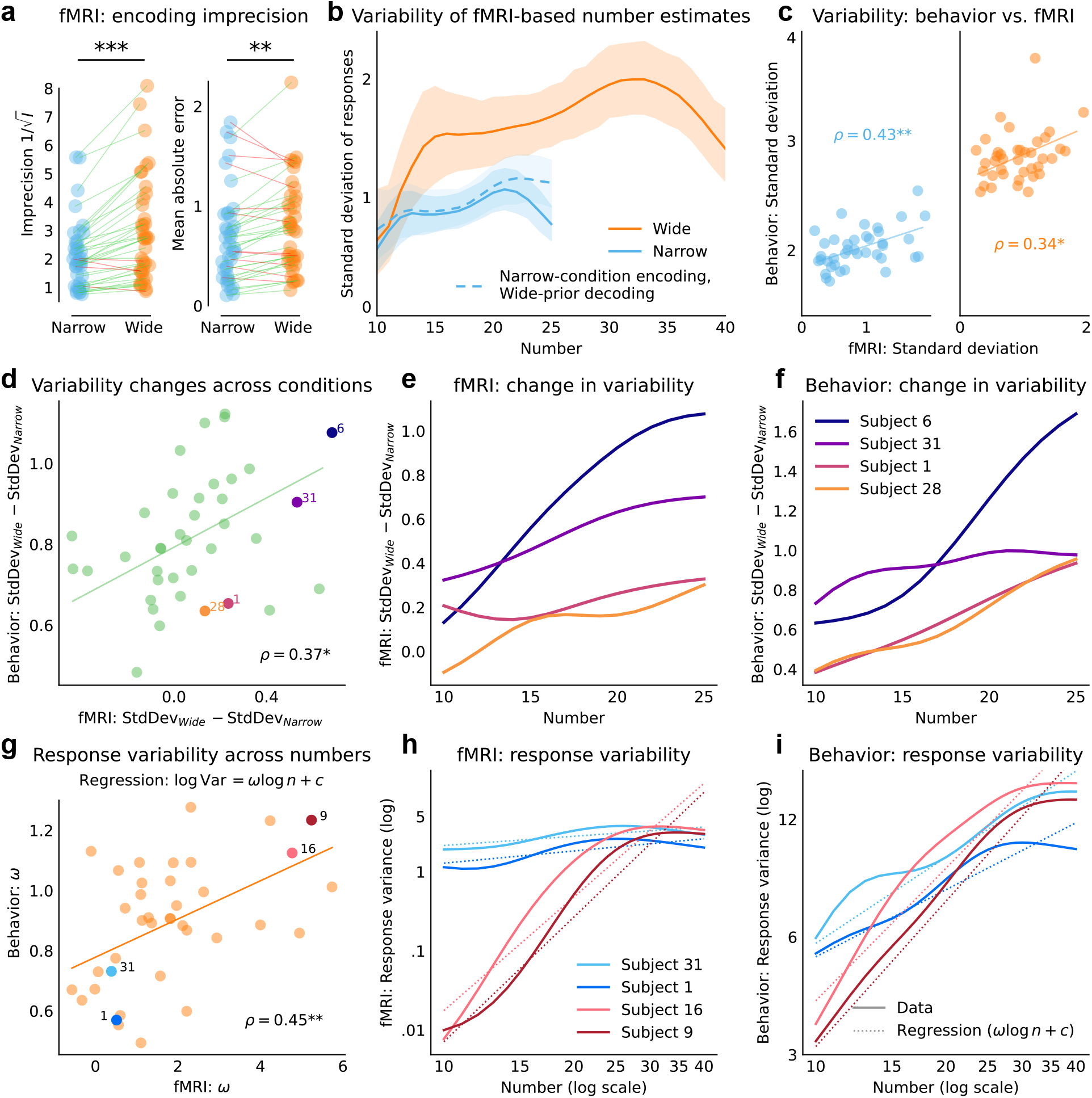
The neural-code imprecision correlates with the behavioral imprecision. **a,** *Left:* fMRI-derived neural imprecision, measured as the inverse of the square-root of the encoding Fisher information, for each participant in each condition. *Right:* Mean absolute error of fMRI-derived estimates. In both cases, a green (red) line indicates an increase (decrease) of the quantity in the Wide condition. **b,** Standard deviation of fMRI-derived numerosity estimates, based on the best-fitting nPRF encoding models in each condition, and on the prior in each condition (solid lines), or on the Wide prior (dashed line), as a function of the presented number. Shaded areas show the 95% confidence intervals. **c,** Standard deviation of the numerosity estimates provided by the participants vs. derived from fMRI data, in the Narrow (left) and Wide (right) conditions (in each panel, one point corresponds to one participant). **d,e,f,** Analysis of the change in imprecision across conditions: **d,** Across-conditions difference in the standard deviation of estimates, as measured in the behavioral data vs. as derived from the fMRI data. A larger increase in fMRI-derived response variability correlates with a larger increase in behavioral response variability. **e,f,** Across-conditions difference in the standard deviation of estimates as a function of the presented number, as derived from the fMRI data (**e**) and from the behavioral data (**f**), for four participants that show a diversity of adaptation profiles across numbers. The four participants are shown in **d** with matching colors. **g,h,i,** Analysis of the imprecision varying across numbers: **g,** Parameter *ω* governing each participant’s degree of compression across numbers in the Wide condition (see regression), estimated from the behavioral data vs. from the fMRI data. **h,i,** Response variance (log scale) as a function of the presented number (solid lines), as derived from the fMRI data (**h**) and from the behavioral data (**i**), for four participants that show a diversity of compression across numbers, and corresponding regression lines (dashed lines). The four participants are shown in **g** with matching colors. Individual data in e,f,h,i were smoothed with Gaussian kernels of widths 0.2 (e,g; log scale) and 3 (h,i) to emphasize trend. Panels d,g omit one outlier with abscissa value 4.7 standard deviations or more away from group mean. ***: *P <* 0.001, **: *P <* 0.01, *: *P <* 0.05.

Specifically, we generate noisy neural responses from the encoding model and decode them into number estimates, by taking the posterior mean. We obtain distributions of such simulated fMRI-derived numerosity estimates, by repeating this procedure 20,000 times for each participant, numerosity, and condition (see also Methods). The mean absolute error of these simulated decoded estimates is significantly larger in the Wide condition than in the Narrow condition (paired t-test: *t*(38) = 3.48, *P* = 0.0013; Fig. 5a, right panel), consistent with the Fisher-information analysis above. Moreover, the standard deviation of the fMRI-derived numerosity estimates as a function of the true numerosities mirrors the pattern of participants’ behavioral variability (compare Fig. 5b to Fig. 1b), with consistently higher variability in the Wide condition.

As a control analysis, we also decode simulated neural responses from the Narrow condition using the Wide prior. This manipulation only modestly increases the variability relative to the Narrow prior, with the main difference being the absence of a boundary effect at 25 (Fig. 5b, dashed line). This result indicates that the increased variability in the Wide condition is primarily driven by changes in neural encoding, rather than by the decoding prior.

A more traditional approach to estimating the acuity of neural representations is to decode empirical fMRI data using a cross-validated scheme^44^. However, we believe such an approach is less effective for quantifying precise measures of uncertainty across the number line, as noise from encoding model estimation and fMRI measurement variability in individual test-set trials would compound (see also Ref. [54]). Still, for the sake of completeness, we applied an established Bayesian inversion method for encoding models^44,50,51,53,55^ to decode the presented numerosity (see Methods). Decoding was reliable, with a repeated measures correlation between decoded and presented numerosity of *r*_rm_(18615) = 0.341, *P* ≈ 0 when both conditions were combined, *r*_rm_(9277) = 0.210, *P* = 4.4 × 10*^−^*^93^ for the Narrow condition, and *r*_rm_(9298) = 0.189, *P* = 2.8 × 10*^−^*^75^ for the Wide condition. Note that the combined correlation is somewhat inflated, as the decoder requires knowledge of the ground-truth condition to select the appropriate encoding model parameters (e.g., preferred numerosity). The within-condition correlations are themselves substantial, however, and consistent with decoding accuracies reported in earlier numerosity studies^44^. Critically, even when decoding in the Wide condition was restricted to trials in which the numerosity fell within the Narrow range (i.e., *n* ≤ 25), with the Wide-condition posterior censored at *n* = 25 before renormalising so that in both conditions the posterior was integrated over the same range, the average decoder error was significantly larger in the Wide condition (mean: 4.96, SD: 0.44) than in the Narrow condition (mean: 4.83, SD: 0.37; one-sided paired *t*-test: *t*(38) = 1.91, *P* = 0.032), confirming reduced encoding precision in the Wide condition even over a matched stimulus range. Moreover, when using the decoded uncertainty (i.e., the standard deviation of the decoded posterior) as a measure, this was significantly larger in the Wide compared to the Narrow condition (mean: 0.78, SD: 0.36 versus mean: 0.66, SD: 0.37; *t*(38) = 3.88, *P* = 2.0 × 10*^−^*^4^). In sum, the decoding analysis corroborates our main findings, confirming that the absolute acuity of the population-level neural representation is reduced in the Wide condition.

## Changes in neural coding correlate with changes in behavioral variability

The fMRI data suggest decreased precision of the neural representations of numbers in the Wide condition (Fig. 5a,b), and the behavioral data exhibit an increase in the participants’ response variability (Fig. 1b). Both findings are consistent with dynamic efficient coding, i.e., a reallocation of representational resources when the range widens, leading to less precise encoding. If the behavioral variability proceeds from the imprecision in neural encoding, then one would expect that these two quantities, and their changes across priors, should be correlated. We thus examine whether the statistics of the fMRI-derived numerosity estimates correlate with those of the participants’ estimates.

Prior studies have shown that participants with less precise tuning in right IPS are more imprecise in binary choices involving numbers^43,44,50^. Consistent with these findings (but here with an estimation task), we find that the standard deviations of behavioral estimates and of fMRI-derived estimates are significantly correlated across participants, in both conditions (Narrow: Spearman’s *ρ* = 0.43, *P* = 0.0059, *N* = 39; Wide: *ρ* = 0.34, *P* = 0.036, *N* = 39; Fig. 5c). In short, less precise neural encoding correlates with less precise behavioral responses.

Turning to the question of adaptation, i.e., to the change in encoding across conditions, we first note that participants exhibit individual differences in the degree to which their neural encoding precision adjusts across priors. To capture these differences, we fit a ‘Participant-specific shifts and width-scaling’ model with two parameters: a slope parameter, *r*_µ_, governing shifts in preferred numerosities (i.e., *µ*_w_ = 10 + *r*_µ_(*µ*_n_ − 10) if *µ*_n_ ≥ 10, otherwise *µ*_w_ = *µ*_n_), and a width-scaling parameter, *r*_σ_, characterizing changes in tuning width (i.e., *σ*_w_ = *r*_σ_*σ*_n_). With this model, we ask whether individual variations in neural adaptation correlate with individual variations in behavioral adaptation. We thus examine the difference across conditions of the standard deviation of responses (StdDev_Wide_ − StdDev_Narrow_). The difference computed with the behavioral estimates is significantly correlated with the difference computed with the fMRI-derived estimates (Fig. 5d; Spearman’s *ρ* = 0.34, *P* = 0.036, *N* = 39.) Thus, participants whose neural populations show larger losses of precision with the Wide prior also show greater increases in behavioral variability. Figure 5e,f illustrates for four representative participants how the behavioral changes in variability parallels the fMRI-derived changes in variability. This close link between shifts in neural coding and behavioral noise is again consistent with our hypothesis that the dynamic shifting of numerosity tuning functions implements dynamic efficient coding.

## Neural and behavioral imprecision across numbers

Finally, participants also differ in how their precision varies across numbers. Number representations are roughly consistent with a logarithmic compression, but there are individual variations^56,57^. Our data allow us to test whether such individual differences are reflected in parietal tuning profiles. We quantify the compression in both neural and behavioral data by fitting a power law relating response variance (Var) and numerosity (*n*), as log Var = *ω* log *n* + *c*. This relationship, which can be understood as a generalization of Weber’s law (obtained with *ω* = 2), provides good fits to both behavioral (mean *R*^2^ = 0.57, IQR: 0.50-0.68) and fMRI-derived estimates (mean *R*^2^ = 0.47, IQR: 0.21-0.74). Here we focus our analysis on the Wide condition, as it encompasses a broader range of numerosities.

Behavioral and neural estimates of *ω* are significantly correlated (Fig. 5g; Spearman’s *ρ* = 0.45, *P* = 0.0047, *N* = 38, when excluding one outlier whose behavioral *ω* is 4.7 standard deviations from the group mean; and *ρ* = 0.40, *P* = 0.011, *N* = 39, when including the outlier). Thus, participants whose neural populations show steeper increases in encoding imprecision across numerosities also exhibit stronger behavioral compression (Fig. 5h,i).

## Discussion

Efficient coding is a well-established principle in low-level sensory systems, and behavioral studies suggest that it underlies the representation of numerosity and subjective value^38–41,58^. Yet it remains unclear how this principle is dynamically implemented by neural populations, including for the encoding of abstract magnitudes used for decision making. Here we show that numerosity-sensitive populations in human parietal cortex adapt their tuning to the contextual statistics of numerical stimuli. Consistent with a dynamic implementation of efficient coding, the encoding precision decreases under the wider prior. The observed adjustments in preferred numerosities and receptive-field widths follow a predictable pattern, supporting the proposal of distributed range adaptation, whereby populations encode stable quantiles across contexts. Moreover, individual differences in these neural adjustments correlate with individual differences in response variability, suggesting that the observed neural activity underlies behavior. In short, the scaling of the prior prompts the scaling of the neural encoding, which in turn underlies the scaling of the behavioral variability.

Our study extends efficient coding in several ways. First, we show in neural data that it applies beyond the perception of physical quantities to abstract magnitudes such as numerosity. Second we show, consistent with proposals based on behavioral data^38–41,58^, that it is a dynamic process which flexibly adapts (on the timescale of a one-hour experiment) to changing statistical contexts. Third, we uncover a neural mechanism, distributed range adaptation, through which tuning properties adjust to optimize encoding for the input distribution.

Distributed range adaptation entails a quantitative prediction about the specific shifts of the neural populations’ preferred numerosities across priors (Eq. 1). In modeling population receptive fields, this theoretical constraint greatly reduces the flexibility of a model, but yields a more robust fit than when unconstrained (as evidenced by the increased cross-validated variance explained). Thus, the hypothesized constraint in fact captures a structuring property of the neural data. Similarly, the receptive-fields widths are well captured by a scaling relation, whereby the wider prior results in wider receptive fields. The best account of the data is obtained by assuming that all the participants shift their preferred numerosities by the same factor, and scale their receptive-field widths by the same factor. This homogeneity further substantiates our conclusions and suggests that distributed range adaptation is a standard computational mechanism that structures how sensory networks reorganize in adaptation to changing contexts.

Individual variations nevertheless remain, and enable us to tie participants’ behavior to their neural activity. Our results support, but also go beyond, the finding that participants with less precise neural encoding are also less precise in behavior^43,44,50^. We show that a participant who adjusts their neural tuning to a greater extent across priors than the average participant also exhibits larger changes in behavioral variability. Similarly, participants whose neural encoding suggests a stronger compression of larger numbers also exhibit stronger diminishing sensitivity for larger numerosities, supporting the hypothesis that Weber’s law originates in the tuning properties of parietal neurons^17^. In other words, whether we look at variations across conditions or variations across numbers, we find that the precision of the neural encoding correlates with the precision of the behavior.

Across priors, the widths of the receptive fields scale by a factor smaller than that applied to the prior (1.3 vs. 2), as conjectured. This implies that, relative to the size of the prior, the specificity of the encoding populations is higher with the Wide prior. This is consistent with a recent efficient-coding model of endogenous precision, which predicts that wider priors should result in lower absolute precision but higher relative precision, so as to mitigate the larger errors incurred under the wider priors^38^.

Our findings were made possible by a combination of methodological choices: we study a relatively large sample of participants (39; with two sessions each), performing an estimation task instead of a (more typical) binary-choice task, and we make extensive use of advanced encoding models of fMRI data, namely, numerical population receptive fields models, to formally quantify the acuity of the neural code. This enables us to effectively test the neural data against targeted hypotheses, and to measure with precision the attributes of the neural substrates that support human representations. In particular, the adaptation mechanism that we propose is consistent with the findings of Kido *et al.*^46^, who use a discriminative multivariate analysis of fMRI data obtained in a numerosity bisection task to show that neural activity patterns contain relative-rank information about numerosities. Our different approach—a generative forward-encoding framework whereby non-linear tuning functions are explicitly modeled—allows us to directly exhibit the tuning properties of encoding populations, and to quantify their adjustments across contexts. Further, this enables us to test, and confirm, our specific hypothesis of distributed range adaptation as the mechanism mediating dynamic efficient coding, and supporting adaptive behavior.

In rats, a pattern similar to distributed range adaptation has been observed in the representation of elapsed time in striatal and hippocampal populations^23,24^, suggesting that it may be a canonical property of sensory networks. The notion of distributed range adaptation moreover seems to partially extend to the encoding of spatial location. When a rat’s environment is expanded, about 36% of its recorded hippocampal place cells exhibit a corresponding rescaling of the locations of their receptive fields, such that they code for the same relative position across environments, consistent with a 2D extension of our results^59,60^. Notably, the areas of the receptive fields of these place cells increase with the area of the environment, but they scale by a ratio smaller than the scaling factor applied to the environment^59^, analogous to our results regarding the receptive-fields widths. This similarity in findings across species and neural areas suggests a similar strategy of increased relative precision to mitigate larger errors in larger spaces.

One limitation of our study is that we did not fully control for all low-level visual features beyond numerosity, such as total surface area or dot density. Controlling for such features inherently introduces new confounds (e.g., when total area is held constant, dot size becomes a secondary cue to numerosity). Correlations of these features with numerosity can be reduced by varying, across trials, how numerosity relates to the physical dimensions of the stimulus, but it is difficult, if not impossible, to eliminate them^61–63^. Our experimental design cannot exclude the hypothesis of such confounds. We note however several reasons to believe that our results reflect genuine numerosity encoding rather than low-level visual processing. First, extensive prior work has demonstrated that numerosity tuning in IPS is robust to variations in low-level visual properties^33,47^. Second, activity in this area has been shown to be consistently better explained by numerosity than by these visual features, suggesting that these parietal representations are indeed about numerosity^64,65^. Third, a direct link has been established between the acuity of numerosity tuning in IPS and decision-making about Arab numerals^44^, further solidifying the connection between numerosity tuning and numerical cognition rather than low-level visual processing. Finally, our focus was not on isolating numerosity as a unique stimulus dimension, but rather on the mechanism of adaptive population coding—how tuning curves reorganize across contexts. Importantly, our theoretical account is robust to this ambiguity: regardless of whether the encoded magnitude is numerosity or a correlated physical dimension, the efficient-coding framework makes the same predictions for the contextual reorganization of tuning curves, and the corresponding behavioral shifts.

A second potential limitation concerns the potential contribution of spatial coding. Posterior parietal cortex encodes retinotopic spatial location^66–68^, and our task required participants to report numerosity via a spatially varying response bar, raising the question of whether some tuning reflects spatial attention or motor planning rather than numerosity per se. Several converging lines of evidence argue against this interpretation. First, a retinotopic account predicts foveal tuning to dominate, corresponding to the center of the response bar and therefore numbers in the middle of the ranges. However, preferred numerosities in our data are left-skewed, directly contradicting this prediction. Second, numerosity preferences in parietal cortex have been found to be independent of receptive field position and size^69,70^, and numerosity-tuned responses are found even in populations whose receptive fields do not overlap with the stimulus. This demonstrates that numerosity tuning in IPS is not a byproduct of retinotopic organization. Third, a spatial confound would predict an additive shift in preferred numerosities between conditions — a fixed offset reflecting a change in response bar position — whereas we observe a multiplicative, slope-like relationship in which the magnitude of the shift scales with preferred numerosity (see Supplementary Figure S17 for a graphical illustration). This pattern is the signature of range adaptation, not spatial remapping.

Distributed range adaptation requires encoding populations to update their tuning properties, and to do so in a mutually consistent manner, to maintain a coherent collective code. Behavioral evidence suggests that this adaptation may occur relatively quickly (on the order of a second^39^). Changes in neuronal tuning could be implemented via synaptic reweighting^71,72^, or via gain adaptation in a recurrent network^39,73^. Network updates could be driven by top-down modulatory signals, or by local learning rules implementing quantile regression^74^, as in recent models of distributional reinforcement learning^75,76^. Thus, while our work establishes that the neural code for number flexibly and efficiently adapts to context, future studies using electrophysiological recordings^31^, neuromodulatory interventions^77^, or layer-specific neuroimaging^78^ should further adjudicate which mechanisms drive this adaptation. These approaches may clarify potential dysfunction in clinical conditions—such as gambling and drug-use disorders—where distorted magnitude representations may contribute to maladaptive decision-making^79,80^. For example, recent opioid use has been shown to bias reinforcement learning toward absolute-value coding rather than range-adapted coding: individuals seem to fail to rescale value representations to the locally relevant reward range, and exhibit decreased performance when this range is narrower^80^. In the framework of distributed range adaptation, this pattern would correspond to a reduced or absent shift and rescaling of population tuning when the value range changes. Such a failure of distributed range adaptation may thus provide a mechanistic account of reduced context adaptation in human addiction.

## Methods

### Task and participants

Thirty-nine healthy participants (13 females, aged 18-34, mean age 23.1) participated in the experiment. Participants were provided information about the objective of the study, the equipment used, the data recorded, the task, as well as the payoff mechanism. Participants were screened for MR compatibility. No participant showed indications of psychiatric or neurological disorders or needed visual correction. The experiment conformed to the Declaration of Helsinki and was approved by the Canton of Zurich’s Ethics Committee. All the participants provided their written consent after being presented with the relevant information.

Each participant completed two experimental sessions, each lasting approximately one hour, with an additional 45 minutes for preparatory procedures such as changing into scanning attire and scanner setup. Each session included two blocks corresponding to the Narrow and Wide conditions, presented in pseudo-random, counterbalanced order.

Each block started with a *learning* phase (15 trials). In each trial of this phase, the participant was shown a stimulus together with its objective numerosity, represented by an Arabic numeral. The participant could progress from one trial to the next at their own pace, and no further response was asked of them in this phase.

The *learning phase* was followed by the *feedback phase* (30 trials), during which participants estimated the numerosity of the presented stimulus and received feedback on the ground truth numerosity after each response. Each trial began with a green fixation cross (300 ms), followed by a red fixation cross (300 ms), both presented on a gray background. The stimulus array then appeared for 600 ms, consisting of 10–40 white dots (0.1*^◦^* visual angle) within a 2.5*^◦^* circular aperture. Both the aperture and background shared the same gray color, thus its edges were not directly visible. Immediately after the presentation of the stimulus array, a slider appeared, scaled proportionally to the current prior range. Participants used an MRI-compatible trackball to indicate their perceived numerosity, with no time limit for their response. After response submission, the correct numerosity was displayed as feedback for 500 ms. The *learning* and *feedback phases* aimed to familiarize participants with the stimuli and the prior distribution of the current condition.

After the *learning* and *feedback* phases, participants performed the *main task* (see Fig. 1a). Each trial in the *main task* started with a green fixation cross (300 ms), followed by a red fixation cross (300 ms), and then the stimulus array, which was presented for 600 ms. Crucially, a variable delay of 4, 5, or 6 seconds (randomly selected) followed the stimulus array, during which only a white fixation cross was displayed. This temporal separation ensured that stimulus-related BOLD activity was not contaminated by decision-making and motor response-related BOLD activity^44,50,51^. In the main task, participants were given three seconds to provide their estimate. Each block of the main task comprised 120 trials (4 runs of 30 trials). Thus, each session included 240 trials, and a total of 480 trials were collected per participant (240 trials per condition).

At the end of the experiment, participants received a reward consisting of a participation fee (30 CHF/hour) plus a performance bonus. The performance bonus was calculated based on estimate accuracy: for each trial, an amount equal to CHF 0.07 −(x^-x)^2^*/*600 was added to the bonus, where x^ is the participant’s estimate and *x* is the ground truth numerosity. Trials from both the *main task* and the *feedback*-phase contributed to the performance bonus. Late responses incurred a penalty of CHF 0.10 per trial, deducted from the bonus. All participants earned a positive performance bonus, with an average of CHF 29.77 (sd: CHF 6.00) across the two sessions (CHF 1 ≈ USD 1.25 at current exchange rates).

### MRI data acquisition

We acquired structural and functional MRI data using a Philips (Best, the Netherlands) Achieva 3T whole-body MR scanner equipped with a 32-channel MR head coil, located at the Laboratory for Social and Neural Systems Research (SNS-Lab) of the UZH Zurich Center for Neuroeconomics. In both sessions, we collected 8 runs of fMRI data with a T2*-weighted gradient-recalled echo-planar imaging (GR-EPI) sequence (132 volumes + 5 dummies; flip angle 90 degrees; TR = 2286 ms, TE = 30ms; matrix size 96 × 70, FOV 240 × 175mm; in-plane resolution of 2.5 mm; 39 slices with thickness of 2.5 mm and a slice gap of 0.5mm; SENSE acceleration in phase-encoding direction (left-right) with factor 1.5; time-of-acquisition 5:02 minutes). At the beginning of each session, we acquired high-resolution T1-weighted 3D MPRAGE image (FOV: 256 × 256 × 170 mm; resolution 1 mm isotropic; Shot TR = 2800 ms; TI = 1098.6 ms; 256 shots, flip angle 8 degrees; TR = 8.3 ms; TE = 3.9 ms; SENSE acceleration in left-right direction 2; time-of-acquisition 5:35 minutes), while participants performed the *learning* and *feedback* phases.

### MRI preprocessing

Preprocessing of fMRI data was performed using *fMRIPrep* 23.2.1^81,82^, which is based on *Nipype* 1.8.6^83,84^.

#### Preprocessing of B0 inhomogeneity mappings

A *B0*-nonuniformity map (or *fieldmap*) was estimated based on two or more echo-planar imaging (EPI) references using topup^85^ (FSL).

#### Anatomical data preprocessing

Each T1w image was corrected for intensity non-uniformity (INU) using N4BiasFieldCorrection^86^, distributed with ANTs 2.5.0. The T1w reference was then skull-stripped using a *Nipype* implementation of the antsBrainExtraction.sh workflow (ANTs), with OASIS30ANTs as the target template. Brain tissue segmentation of cerebrospinal fluid (CSF), white matter (WM), and gray matter (GM) was performed on the brain-extracted T1w using fast^87^ (FSL). An anatomical T1w-reference map was computed after registration of the 4 INU-corrected T1w images using mri_robust_template^88^ (FreeSurfer 7.3.2). Brain surfaces were reconstructed using recon-all^89^ (FreeSurfer 7.3.2). The brain mask estimated previously was refined using a custom variation of the method to reconcile ANTs-derived and FreeSurfer-derived segmentations of the cortical gray matter^90^. Volume-based spatial normalization to the MNI152NLin2009cAsym standard space was performed through non-linear registration with antsRegistration (ANTs 2.5.0), using brain-extracted versions of both the T1w reference and the T1w template. The following template was selected for spatial normalization and accessed via *TemplateFlow*^91^ (version 23.1.0): *ICBM 152 Nonlinear Asymmetrical template version 2009c*^92^ (TemplateFlow ID: MNI152NLin2009cAsym).

#### Functional data preprocessing

For each of the 16 BOLD runs per participant (across all tasks and sessions), the following preprocessing steps were performed. First, a reference volume was generated using a custom *fMRIPrep* methodology for head-motion correction. Head-motion parameters (transformation matrices and six corresponding rotation and translation parameters) were estimated with respect to the BOLD reference using mcflirt^93^ (FSL), prior to any spatiotemporal filtering. The estimated *fieldmap* was aligned to the target EPI reference run using rigid registration, and the field coefficients were mapped onto the reference EPI.

The BOLD reference was co-registered to the T1w reference using bbregister (FreeSurfer), which implements boundary-based registration^94^. Co-registration was configured with six degrees of freedom. Several confounding time series were calculated based on the preprocessed BOLD data: framewise displacement (FD), DVARS, and three region-wise global signals (CSF, WM, and whole-brain). FD was computed using two formulations: absolute sum of relative motions^95^ and relative root mean square displacement between affines^93^. FD and DVARS were calculated for each functional run using their *Nipype* implementations^95^. Many internal operations in *fMRIPrep* use *Nilearn* 0.10.2^96^. For further details, see the *fMRIPrep* workflow documentation.

### fMRI analysis

The key aim of our fMRI analyses was to estimate the nonlinear tuning of neuronal populations in the parietal cortex to numerosity, and how this tuning was shaped by different task contexts (i.e., priors), following established approaches^33,44,50,51^. After preprocessing, the analysis proceeded in three phases:

1. Estimating voxelwise BOLD responses for every trial using a single-trial GLM.
2. Fitting a large set of numerical population receptive field (nPRF) models, differing in complexity and scientific assumptions, and comparing them using cross-validated *R*^2^.
3. Using statistical measures to quantify the information about numerosity contained in parietal activity, and how this varied with task context and numerosity.

These phases will be described in detail in the following sections.

#### Single-trial BOLD response estimation

We used the GLMSingle Python package^97^ to estimate single-trial BOLD response amplitudes. This package employs cross-validation to optimize a generalized linear model (GLM) by selecting: 1) the most suitable hemodynamic response function from a predefined library, 2) the optimal L2-regularization parameter to mitigate the effects of correlated single-trial regressors, and 3) GLMDenoise regressors^98^, derived from the first *n* principal components of noise voxels (defined as voxels with low explained variance in the task-based GLM). The number of principal components was determined via cross-validation.

To construct the design matrix for GLMSingle, we modeled the onsets of both (1) the stimulus arrays of different numerosities and (2) the response periods. Trials were categorized for cross-validation as either stimulus trials (e.g., stimulus_10) or response trials (e.g., response_11 or no_response). Based on extensive pilot analyses, as well as earlier studies^50,51^, we chose not to include additional confound regressors (e.g., motion parameters, RETROICOR, or aCompCor) beyond those provided by GLMDenoise. This decision follows prior work showing that additional regressors often correlate with those derived by GLMDenoise and typically do not further improve decoding accuracy; in fact, they can sometimes reduce it by introducing overfitting^50,97,98^.

### nPRF estimation

To estimate nPRF parameters and test our hypotheses , we fitted a number of numerical population receptive field (nPRF) models. These models only differed in the extent to which parameters were (a) constant over conditions (b) completely free between conditions or (c) a linear function of each other (e.g., *µ*_w_ = 10 + *β*(*µ*_n_ − 10)), where *β* could either be fixed or estimated. All nPRF models describes the average responses of a voxel to different numerosities as a function of the number *x*, as follows:

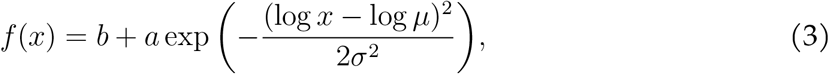

where *b* is the baseline, *a* the amplitude, *µ* the voxel’s preferred numerosity, and *σ* the width parameter of the receptive field.

The nPRF model fitting started with a grid search, making relative activation predictions for each trial using 41 possible preferred numerosities (5-41) and 30 possible width (2-30 in natural space). Grid search is standard in both visuospatial and numerical PRF modeling^33,44,55,99^ as an initial step for continuous optimization, as it is far less likely to converge on local minima. For each voxel, the parameters corresponding to the predicted activation profile with the highest correlation with the actual activation profile was chosen, after which the amplitude and baseline parameters were estimated using ordinary least-squares (OLS). After this initial step, parameters were further optimized using the ADAM gradient descent (GD) optimisation algorithm^100^, as implemented in Tensorflow^101^, with the negative relative explained variance (*R*^2^) as a cost function. Importantly, during this GD optimization, all parameters were jointly fitted, allowing them to move beyond the initial estimates provided by the grid search and OLS.

Some of the models we used put ‘hierarchical‘ constraints on the parameters using scaling parameters *β*. For example, the preferred numerosity of a voxel in the Wide condition was a linear function of the numerosity in the Narrow condition with a slope parameter that is estimated but kept constant across voxels. For these models, the grid search was performed on all voxels, and the mean slope was used as a starting point for the gradient descent optimization.

The nPRF model was fitted both to all voxels within the brain (for visualization purposes), or specifically to voxels in the right NPC. The right NPC mask was taken directly from Barretto-García et al.^44^. In the case of models with hierarchical parameters, this latter approach was necessary, since we did not want voxels outside of the ROI to influence the slope parmameter *β*.

For all models, we fitted both a version of the model in which the average BOLD activity is a function of the *actual ground-truth numerosity* at each trial, and a version in which the BOLD response would be a function of the *response provided by the participant*. The variants that uses the responses generally yielded a higher proportion of variance explained *cvR*^2^ (see Supplementary Information). Thus we use these models for the analyses of the voxels’ tuning (Figs. 3,4). However, when we related the relative acuity of the neural responses to behavioral differences (Fig. 5), we resorted to the models fitted on the ground-truth numerosities. We did so to make sure the fMRI noise estimates and the behavioral responses were statistically completely independent, and avoid ‘double-dipping’.

To identify voxels with reliable numerosity-tuned responses and to facilitate model comparison, we employed an 8-fold cross-validation procedure. For each participant and model, we generated eight distinct train–test splits by iteratively leaving out the *n*th run from both the first and second sessions. This ensured equal representation of the Narrow and Wide conditions in both the training set (210 out of 420 trials each) and the test set (30 out of 60 trials each). The nPRF model was fitted to the training data, and its performance was evaluated using the explained variance (*R*^2^) on the held-out test trials, yielding a cross-validated *R*^2^ (*cvR*^2^). This procedure enabled us to identify voxels with robust and generalizable numerosity tuning while excluding unreliable or noisy responses^52^. Throughout the manuscript, analyses of pRF parameters are restricted to voxels with an average *cvR*^2^ across test runs greater than 0.0.

For model selection at the voxel level, we compared cross-validated explained variance (*cvR*^2^) across candidate models. For each voxel, the model with the highest average *cvR*^2^ across folds was designated as the best-fitting model. Because *cvR*^2^ is evaluated on independent test data, it penalizes overly complex models that overfit noise in the training set. This makes *cvR*^2^ a principled criterion for voxel-wise model comparison, as it favors models that capture reliable neural response mechanisms rather than spurious or idiosyncratic patterns.

For model selection across voxels (e.g., Fig. 3f), we used the proportion of voxels with *cvR*^2^ *>* 0.0. This approach was motivated by the observation that, in noise voxels, nPRF models tend to overfit and can perform much worse than a simple intercept model, yielding very negative (average) *cvR*^2^ values. Consequently, the distribution of *cvR*^2^ across voxels is highly variable with a heavy negative tail. Using the proportion of voxels with positive *cvR*^2^ provides a more stable and interpretable summary measure for model comparison at the population level.

For visualization, voxelwise nPRF parameters were projected onto the cortical surface using the FreeSurfer surfaces generated by fMRIPrep. In particular, we visualized the preferred numerosity of individual vertices. To sample voxelwise pRF parameters onto the surface, we used Nilearn’s vol_to_surf function^96^. Vertices without reliable numerosity tuning were excluded using the *cvR*^2^ *>* 0.0 mask.

All nPRF analyses were implemented using the braincoder package^55^, which provides flexible tools for specifying encoding models and estimating them with a GPU-accelerated TensorFlow backend.

#### Parameter recovery analysis

To verify that our nPRF parameter estimates are not systematically biased by the range of numerosities presented in each condition, we performed a model recovery analysis. We simulated 250 voxels with realistic noise levels, drawing preferred numerosities from the empirical distribution of signal voxels across participants. We then implemented a 2 × 2 factorial simulation design. The first factor was the generating slope relating *µ*_wide_ to *µ*_narrow_, set to either *r*_µ_ = 1 (no shift) or 2 (full range adaptation). The second factor was whether the simulated stimulus design matched our actual experimental design (‘full’) or was restricted to the numerosity range shared between both conditions, i.e., [10, 25] (‘censored’). For each of the four cells, we repeated the simulation 100 times and fit the same constrained nPRF model used in the main analyses. In all cases, the recovered slope parameter was close to the generating value, with negligible bias across both the full and censored designs, confirming that our model fitting procedure is not systematically distorted by differences in stimulus range between conditions. We conducted a similar analysis for the width-scaling parameter, *r*_σ_, and found that it can also be recovered reliably. See Supplementary Information.

#### fMRI-derived Fisher information

To quantify differences in the acuity with which right NPC represents numerosity across participants, conditions, and numerosities, we used the statistical measure of *Fisher Information*^54^. Fisher Information quantifies the sensitivity of a likelihood function to changes in a parameter *θ* (in our case, numerosity *n*), and is defined as

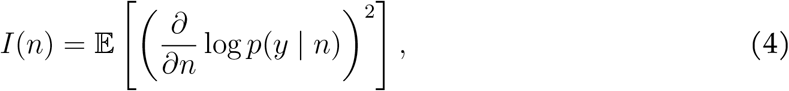

where *p*(*y* | *n*) denotes the likelihood of observing a BOLD activation pattern *y* given numerosity *n*, and the expectation is taken with respect to *p*(*y* | *n*). The inverse of the Fisher Information provides a lower bound on the variance of any unbiased decoder of *n* (the Cramér–Rao bound). Thus, higher Fisher Information implies greater representational acuity and more precise decoding of numerosity.

Thus, our first step was to define a likelihood function *p*(*y n*), following the approach used in our earlier work on decoding numerosities^44,50,51^. The crucial step was to extend the *deterministic* nPRF model,

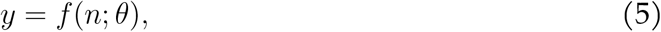

where *θ* = {*µ, σ, A, b*} denotes the preferred numerosity, tuning width, response amplitude, and baseline. This formulation predicts a fixed response pattern for each numerosity, but does not capture trial-to-trial variability in BOLD responses. We therefore extended it into a stochastic model,

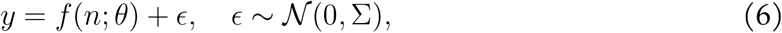

with the noise term *ϵ* accounting for variability in the observed responses.

We estimated Σ by fitting a multivariate normal distribution to the residuals of the best-fitting nPRF model^44,50,51^, using only signal voxels with *cvR*^2^ *>* 0. While relative covariance in noise across voxels is critical for decoding uncertainty^53^, reliable estimation is extremely challenging in a low-*n* regime (480 trials) with hundreds of voxels^102^. Although regularized approaches have been proposed^53^, in our initial analyses these performed worse than assuming diagonal (i.e., uncorrelated) noise. In particular, the decoded posteriors were excessively broad and their decoding performance was worse than a model assuming diagonal noise.

Importantly, our aim here was not to decode trial-by-trial uncertainty (as in^53^ and^50^), but rather to assess how the acuity of numerosity representations varied across participants and conditions. For this purpose, we thus adopted a *diagonal* covariance matrix, assuming no correlations between voxels. Note that these analyses therefore specifically targeted the effects of the *preferred numerosity* and *width* of numerical pRFs, rather than their noise covariance structure. Notably, preliminary analyses revealed no evidence that noise patterns differed between the Narrow and Wide conditions.

We computed Fisher Information for arbitrary neural encoding models using the braincoder package^55^. Since the expectation in Equation 4 cannot readily be expressed in closed form for our models, we evaluated it numerically by sampling from the noise distribution *p*(*y* | *n*). For each sample, we computed the derivative of the log-likelihood with respect to numerosity,

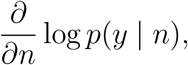

using automatic differentiation. Squaring this term and averaging across samples yields a Monte Carlo estimate of Fisher Information. In practice, we drew 1000 samples for each *n*, providing a stable numerical evaluation of representational acuity at that numerosity. This procedure yielded an estimate of Fisher Information for each numerosity *n*, separately for each participant and condition. These values quantify the acuity of the neural representation at different numerosities. We then used these measures to assess how manipulating the objective prior in the experiment altered the fidelity with which right NPC encoded numerosity across participants and conditions.

#### Expected Variability from an Ideal Bayesian Decoder

Fisher Information is a purely local measure: it reflects sensitivity to infinitesimal changes around a given *n*, but does not account for the discrete nature of number, boundary effects, or the impact of substantial noise^103^. Indeed, in our data, Fisher Information failed to capture the increased precision at the edges of the tested numerosity range. To obtain a more realistic estimate of the expected variability of an actual neural decoder, we therefore complemented the Fisher Information analysis with simulations of an ideal Bayesian decoder. This approach allowed us to approximate the performance of a perfect decoder given the observed neural code, providing a more global measure of representational acuity.

We modeled voxel responses as a multivariate Gaussian distribution,

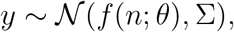

with mean given by the predicted nPRF responses *f* (*n*; *θ*) and covariance Σ estimated from the residuals of the best-fitting model. Using this generative model, we simulated large numbers of single-trial response patterns for each numerosity under both the Narrow and Wide stimulus ranges. For each simulated response *y*, we computed the Bayesian posterior over numerosities, *p*(*n* | *y*), and decoded the stimulus as the posterior mean,

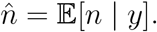

Repeating this procedure many times yielded distributions of decoded numerosities for each true numerosity, from which we quantified variability, systematic bias, and absolute error. By correlating the variability of numerosities decoded from simulated data with the variability observed in participants’ responses, we were able to test whether the precision of neural representations in right NPC correlates with the precision of behavior.

### Decoding analysis

To validate the encoding model fits, we additionally decoded the presented numerosity from single-trial fMRI responses using Bayesian inversion of the encoding model^44,53,55^. For each held-out trial, the likelihood of the observed brain response was computed under the fitted noise model (a Student’s *t* distribution with spherical covariance) for each candidate numerosity value, yielding a posterior probability density function (PDF) over the number line. The posterior mean served as the decoded numerosity estimate for that trial. Voxel selection for decoding was performed via nested cross-validation: within each fold, only voxels that had positive cv*R*^2^ (where the cross-validation of the encoding model was nested within the cross-validation of the decoding model) were included in the decoding of the hold-out test set.

To compare decoding precision between the Narrow and Wide conditions over a matched stimulus range, we restricted the analysis to Wide-condition trials where the presented numerosity fell within the Narrow range (*n* ≤ 25). Crucially, we also censored the Wide-condition posterior PDF at *n* = 25 before renormalising and computing the posterior mean; this ensures that both conditions integrate over the same range and that decoding errors are not inflated by probability mass assigned to values outside the Narrow range. Decoding reliability was quantified with repeated-measures correlation^104^, *r*_rm_, between decoded and presented numerosity. Condition differences in mean absolute error (MAE) were tested with a paired *t*-test across participants.

### Statistical analysis of the behavioral data

Figure 1b shows posterior mean estimates of the group-level parameters from a hierarchical model that includes participant-level random effects. For each condition, the model is defined by the following equations, where *x*^_si_ denotes the estimate of participant *s* in trial *i*, while *x*_i_ denotes the correct number:

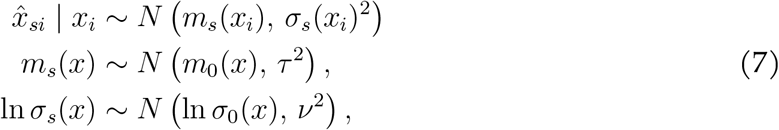

with the priors

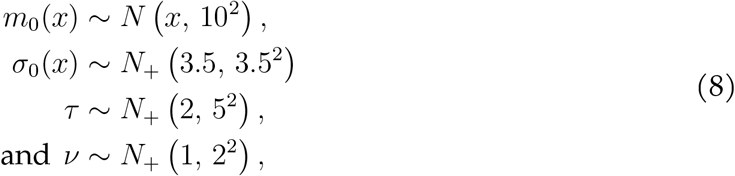

where *N*_+_ is the Gaussian distribution truncated to the positive numbers. This model was estimated using Stan^105^ (10 chains of 1000 samples each, following 1000 warmup iterations, with the HMC-NUTS sampler). The shaded areas in Figure 1b show the 5th and 95th percentile of the posterior.

### Analysis code and data

All analysis code is available on GitHub at https://github.com/Gilles86/neural_priors. The fMRI data will be made publicly accessible on OpenNeuro. Meanwhile, requests can be addressed to the authors.

## Acknowledgements

We are grateful to Cornelia Schnyder and Elena Boldin at the Zurich Center for Neuroeconomics for their excellent assistance with recruitment and participant facilitation and thank Irini Paschalis for scanning support during data collection. C.C.R. received funding from the University Research Priority Program Adaptive Brain Circuits in Development and Learning (URPP AdaBD) at the University of Zurich, as well as from the Swiss National Science Foundation (SNSF, grant no. 100019L-173248). G.d.H. was supported by the Dutch Research Council (NWO, Rubicon grant no. 019.183SG.017/8O3B) and the University of Zurich (Forschungskredit grant no. K-33153-02-01), as well as the URPP AdaBD. SB was supported by a PhD scholarship from the Marlene Porsche Graduate School of Neuroeconomics and funding by the URPP AdaBD. S.J.G. received funding from the U.S. National Science Foundation (NSF, grant no. DRL-2024462), the United States Air Force Office of Scientific Research (grant no. FA9550-20-1-0413), and the Kempner Institute for the Study of Natural and Artificial Intelligence.

## Contributions

A.P.C.: Conceptualization, Data curation, Formal analysis, Writing – original draft, Writing – review & editing. G.d.H.: Conceptualization, Data curation, Formal analysis, Writing – original draft, Writing – review & editing. S.B.: Data curation, Writing – review & editing. S.J.G.: Conceptualization, Writing – review & editing. C.C.R.: Conceptualization, Writing – review & editing.

