## Supplementary Information for "Distributed range adaptation in human parietal encoding of numbers"

### Distributed range adaptation in human parietal encoding of numbers
#### Supplementary Information

Arthur Prat-Carrabin[1,⋆], Gilles de Hollander[2,3⋆], Saurabh Bedi[2,3], Samuel J. Gershman[1], and Christian C. Ruff[2,3]

[1]Department of Psychology and Center for Brain Science, Harvard University, Cambridge, MA, USA
[2]Zurich Center for Neuroeconomics, Department of Economics, University of Zurich, Switzerland
[3]University Research Priority Program (URPP), Adaptive Brain Circuits in Development and Learning, University of Zurich, Zurich, Switzerland

⋆equal contribution
⋆**

August 19, 2026

### 1 Qualitative assessment of nPRF fits

Numerosity-selective population receptive fields (nPRFs) are expected to be considerably noisier than standard visuospatial PRFs, for two reasons: numerosity tuning is much less pronounced in terms of signal-to-noise ratio than retinotopic tuning, and we used an event-related rather than a block design (as is common in visuospatial PRF studies). To nonetheless provide a qualitative sense of model fit quality, we randomly selected 12 voxels from the top 60 voxels ranked by normalised percent signal change (NPCr) with non-monotonic tuning in three example participants (S12, S20, and S26). For each voxel, we plot the mean BOLD response to each numerosity (averaged over $\sim$10 trials) alongside the best-fitting model prediction (efficient shift and fixed increase in dispersion).

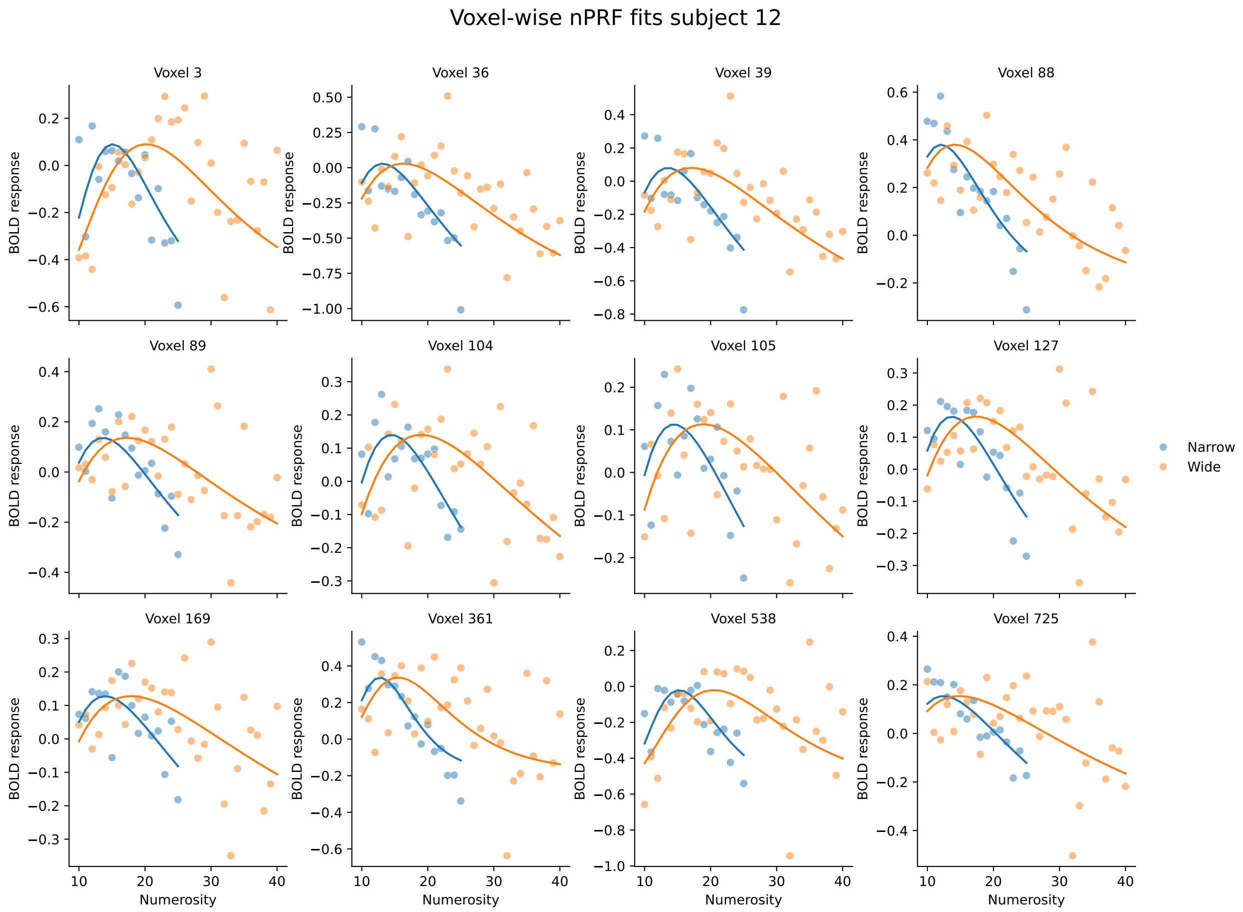


**Fig. S1: Qualitative nPRF model fits for participant 12.** Each panel shows the mean BOLD response (filled circles) as a function of presented numerosity, separately for the Narrow (blue) and Wide (orange) conditions, alongside the fitted nPRF model prediction (solid lines). Voxels were selected from the top 60 NPCr voxels with non-monotonic tuning.

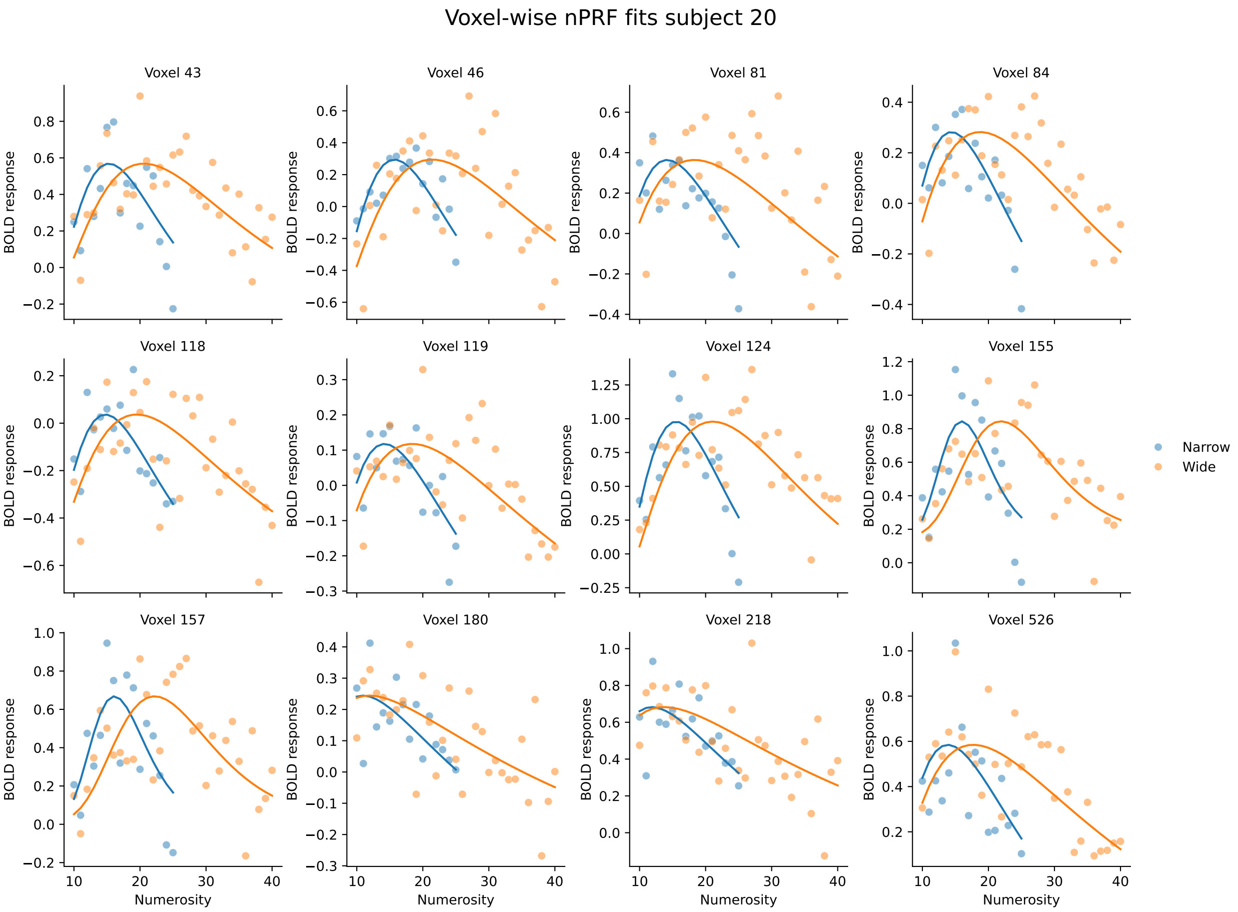


**Fig. S2: Qualitative nPRF model fits for participant 20.** As Fig. S1, for participant 20.

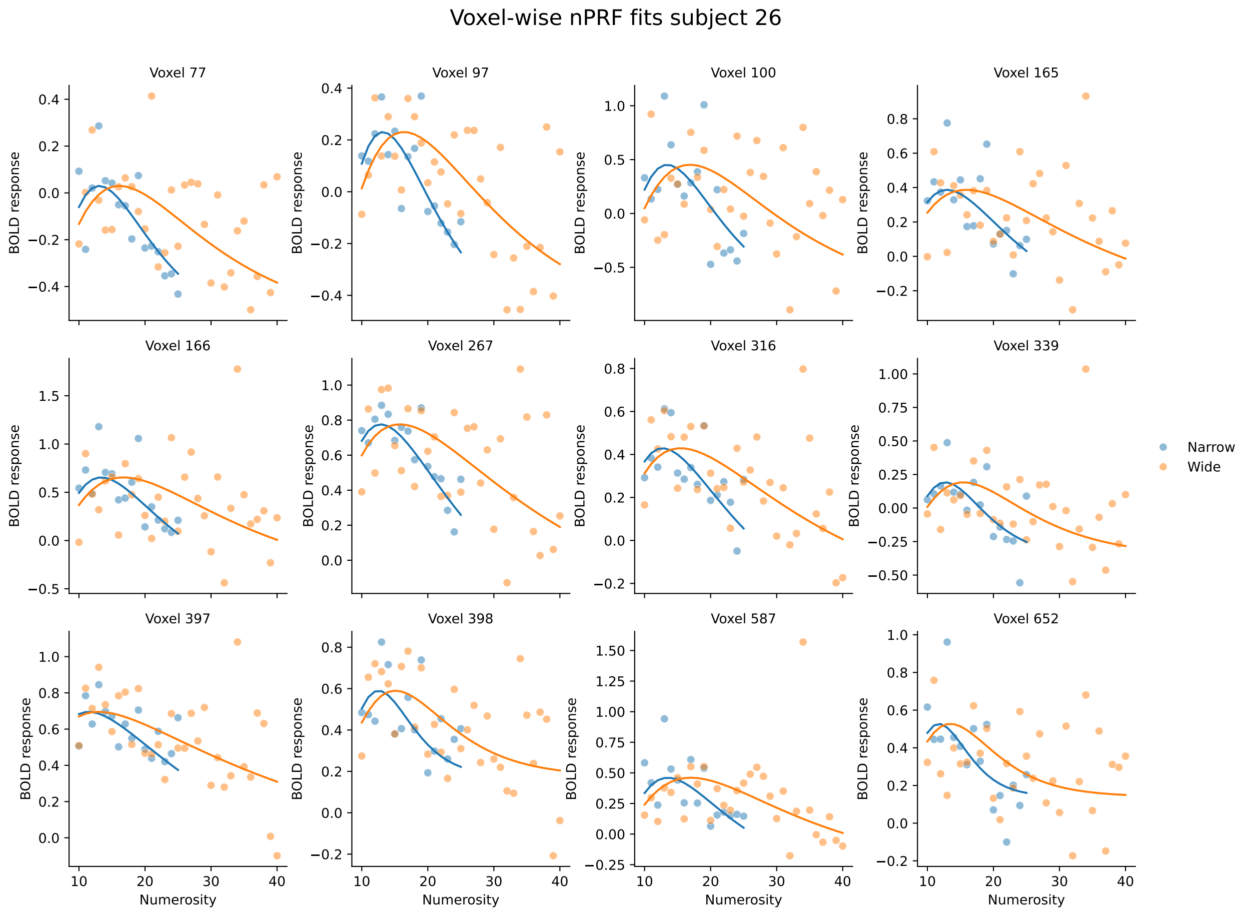


**Fig. S3: Qualitative nPRF model fits for participant 26.** As Fig. S1, for participant 26.

#### 2 Model comparisons

Figure S4 shows the proportion of voxels with cv$R^2 > 0$ with the different nPRF models that we fit. Models fit on the participants' responses (orange line) consistently outperform the models fit on the ground-truth, presented number (teal line).

##### Models with amplitude changes

The precision of the neural encoding depends on the amplitude of the populations' receptive fields, in addition to their preferred numerosities and their widths. Here we test for changes in the amplitudes. Thus we fit models in which the amplitude of each voxel may vary across conditions. Specifically, first we consider a model that features efficient shifts of the preferred numerosities, fixed scaling of the widths (as in the main text), and in which for each voxel we let the amplitudes in the two conditions be free parameters, $a_n$ and $a_w$. This model yields a significantly lower fit than our best-fitting model (with estimates: $t(38) = 6.12$, $P = 3.9 \times 10^{-7}$; with correct numbers: $t(38) = 6.41$, $P = 1.6 \times 10^{-7}$; Fig. S4, model labeled 'Eff. shifts, fixed width-scaling, free amplitudes'). Second, we enforce a relation of proportionality between each participant's voxels amplitudes in the Narrow condition and in the Wide condition, with a fixed ratio for each participant (i.e., $a_w = r_a a_n$). This model yields a better fit than the previous one, but a significantly worse one than our best-fitting model (with estimates: $t(38) = 2.36$, $P = 0.024$; with correct numbers: $t(38) = 2.71$, $P = 0.0099$; Fig. S4, model labeled 'Eff. shifts, fixed width-scaling, p.-sp. amplitude-scaling').

We also consider a 'Participant-specific amplitude-scaling' model, which allows for scaled changes in the amplitudes, while keeping the other parameters constant (rightmost model in Fig. S4). Inspection of all models performance shows that this model does not provide a better account of the data than the efficient-shift model; rather, the evidence points in the opposite direction. With the response-based fit, the 'Participant-specific shifts' model ($\mu_w = 10 + r_\mu(\mu_n - 10)$, with other parameters constant; fifth model) yields a numerically higher proportion of voxels with positive cv$R^2$ than the 'Participant-specific amplitude-scaling' model, and it fits significantly better than the 'No change' model (in which all the parameters are kept identical across conditions; $t(38) = 2.33$, $P = 0.025$). The same result obtains when considering the median cv$R^2$ (see below). By contrast, the 'Participant-specific amplitude-scaling' model does not fit better than the 'No change' model ($t(38) = 1.51$, $P = 0.14$), and it performs significantly worse than the 'Efficient shifts' model (with constant width and amplitude; $t(38) = 2.3$, $P = 0.027$). Thus, allowing for shifts in preferred numerosities offers a better account of the data than allowing for an increase in amplitudes.

##### Models with alternative shift hypotheses

We consider alternatives to our efficient-shift hypothesis (which posits that $\mu_w = 10 + 2(\mu_n - 10)$ if $\mu_n > 10$, otherwise $\mu_w = \mu_n$). First we consider a model in which for all voxels the preferred numerosity in the Wide condition is twice the preferred numerosity in the Narrow condition (i.e., $\mu_w = 2\mu_n$). We compare this model to the model with efficient shifts

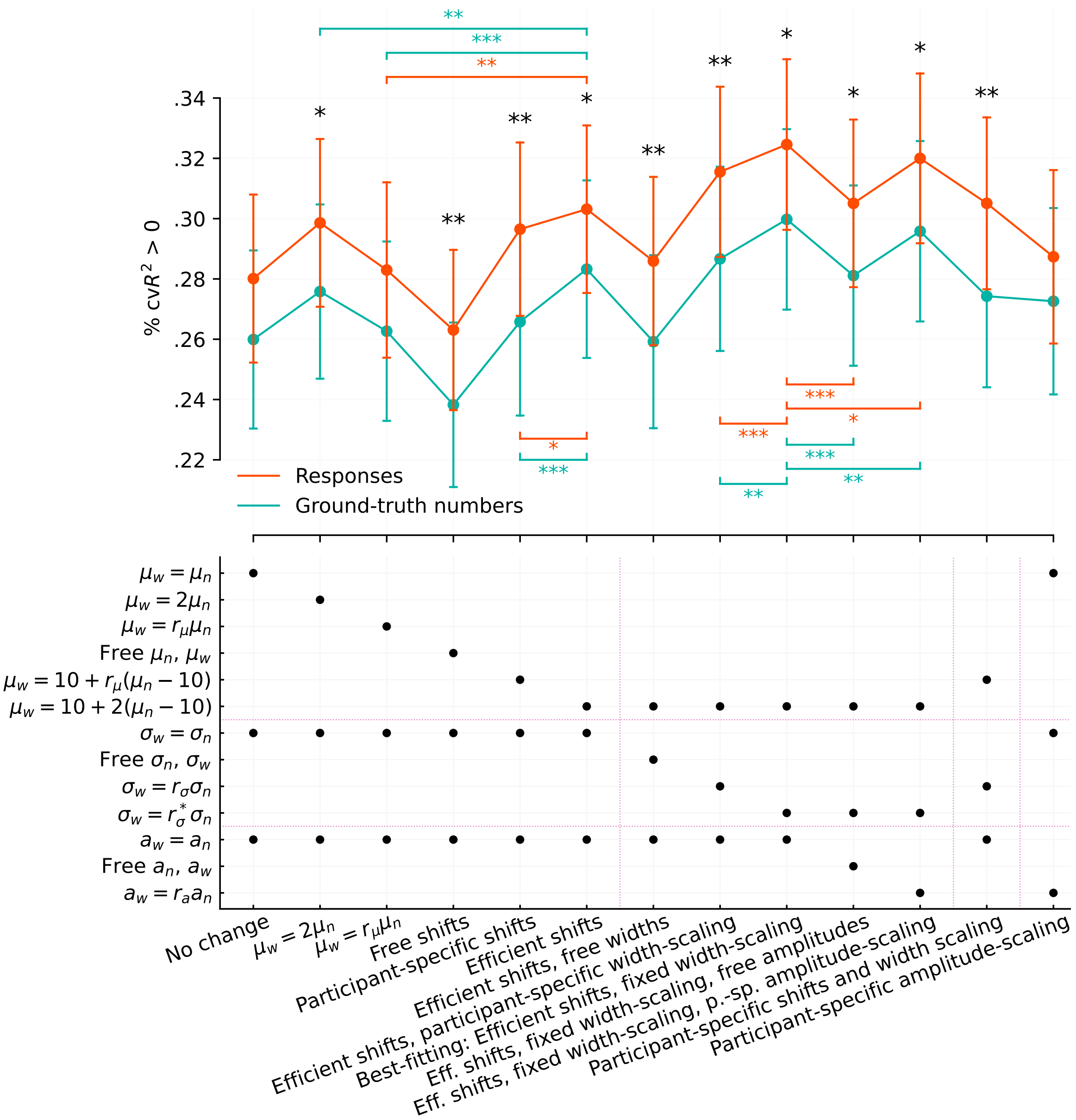


**Fig. S4: Proportion of voxels with positive cross-validated variance explained (cvR$^2$>0) for the different models.** Error bars show $\pm 1$ standard error of the mean. The bottom panel indicates the specifications of the models' parameters across priors. Each model (vertical line) has three dots corresponding to the model's specifications regarding the preferred-numerosity parameters $(\mu_n, \mu_w)$, the width parameters $(\sigma_n, \sigma_w)$, and the amplitude parameters $(a_n, a_w)$, indicated on the left. The ratios $r_\mu$, $r_\sigma$, and $r_a$ are participant-level parameters, while $r_\sigma^*$ is fixed at 1.3 (and shared by all participants). "p.-sp.": participant-specific. ***: $P < 0.001$, **: $P < 0.01$, *: $P < 0.05$.

(and unchanging widths and amplitudes): it yields a lower proportion of voxels with cv$R^2$ positive, with a significant difference when fitting on the correct numbers ($t(38) = 2.63$, $P = 0.012$; Fig. S4, second model). Then we consider a model in which the preferred numerosity in the Wide condition is proportional to that in the Narrow condition, with a fixed parameters for each participant (i.e., $\mu_w = r_\mu \mu_n$; Fig. S4, third model). This model also fits significantly worse than the model featuring the efficient-shift hypothesis (fitting on estimates: $t(38) = 3.19$, $P = 0.003$, fitting on correct numbers: $t(38) = 3.85$, $P = 0.0004$). Thus we conclude that the efficient-shift hypothesis ($\mu_w = 10 + 2(\mu_n - 10)$ if $\mu_n > 10$) better accounts for the data than the hypothesis that $\mu_w$ is proportional to $\mu_n$.

#### Robustness check

To investigate whether our results are robust to the metric used for model comparison, we consider another measure of fit, the median cv$R^2$. The majority of voxels in the ROI do not contain information about the presented numbers and yield very large negative cv$R^2$s. We thus compute for each model the median cv$R^2$ over all the voxels for which at least one model yields a positive cv$R^2$. Figure S5 shows the results obtained with this metric. The figure is very similar to that obtained when looking at the proportion of voxels with positive cv$R^2$ (Fig. S4), and calls for the same conclusions. In particular, the ordering of models is identical, and thus the best-fitting model with this analysis is also the 'Efficient-shifts, fixed width-scaling' model.

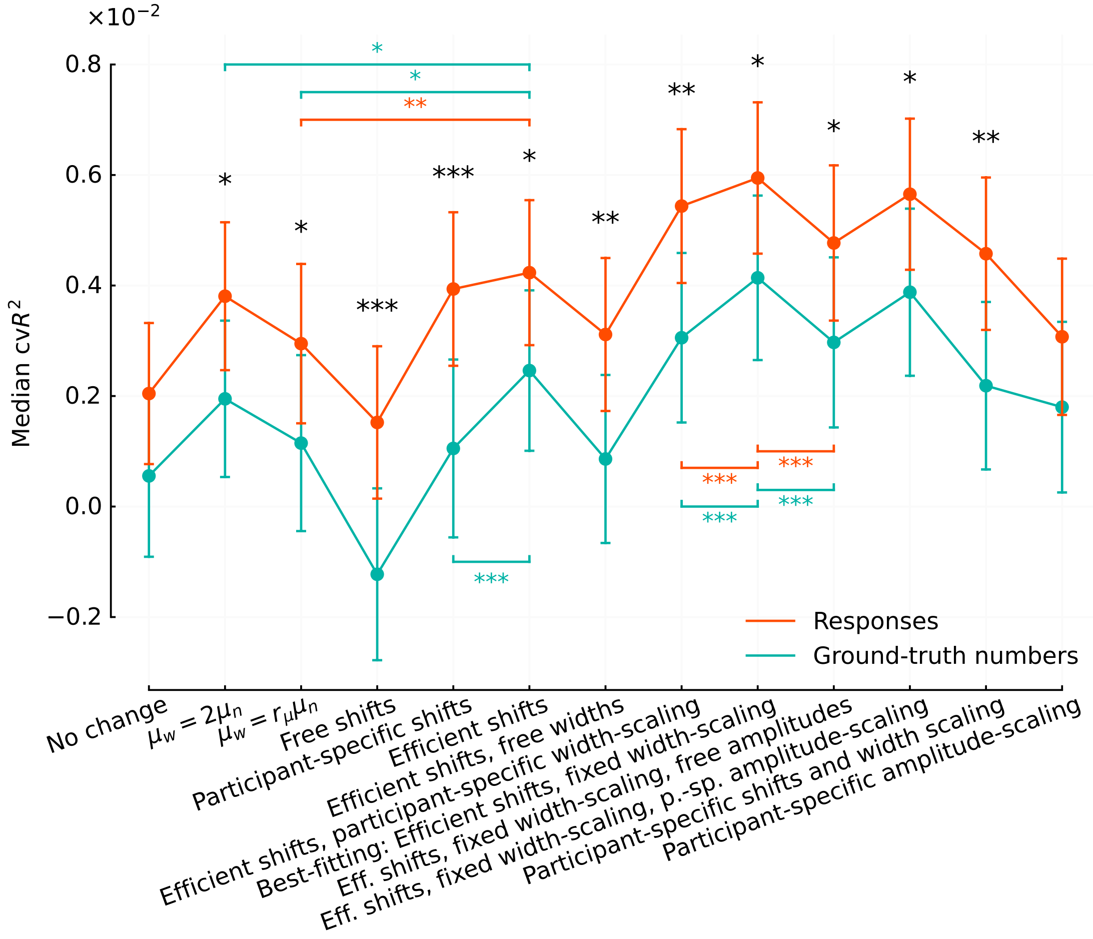


**Fig. S5: Median cvR$^2$ for the different models.** Error bars show $\pm 1$ standard error of the mean. ***: $P < 0.001$, **: $P < 0.01$, *: $P < 0.05$.

### 3 Results when fitting nPRFs with ground-truth numbers

We report the results obtained when fitting the models of population receptive-field with the ground-truth, presented numbers rather than with the participants' responses. Figure S6 is the counterpart of Figure 3 in the main text, and shows the shifts in preferred numerosities across priors. Figure S7 is the counterpart of Figure 4 in the main text, and shows the changes in receptive fields widths across priors. Below, we report for the models fit with the ground-truth numbers the same statistics reported in the main text for the models fit with the responses.

**Preferred numerosities**

- Correlation between preferred numerosities $\mu_n$ and $\mu_w$ in the two conditions: participants pooled: Spearman's $\rho = 0.74$, $P < 10^{-320}, N = 7264$; across participants: average $\rho = 0.64$, interquartile range (IQR): 0.57-0.82.
- Across-participant t-test that the mean shift $\mu_w - \mu_n$ is zero, for $\mu_n \geq 10$: $t(38) = 3.45$,

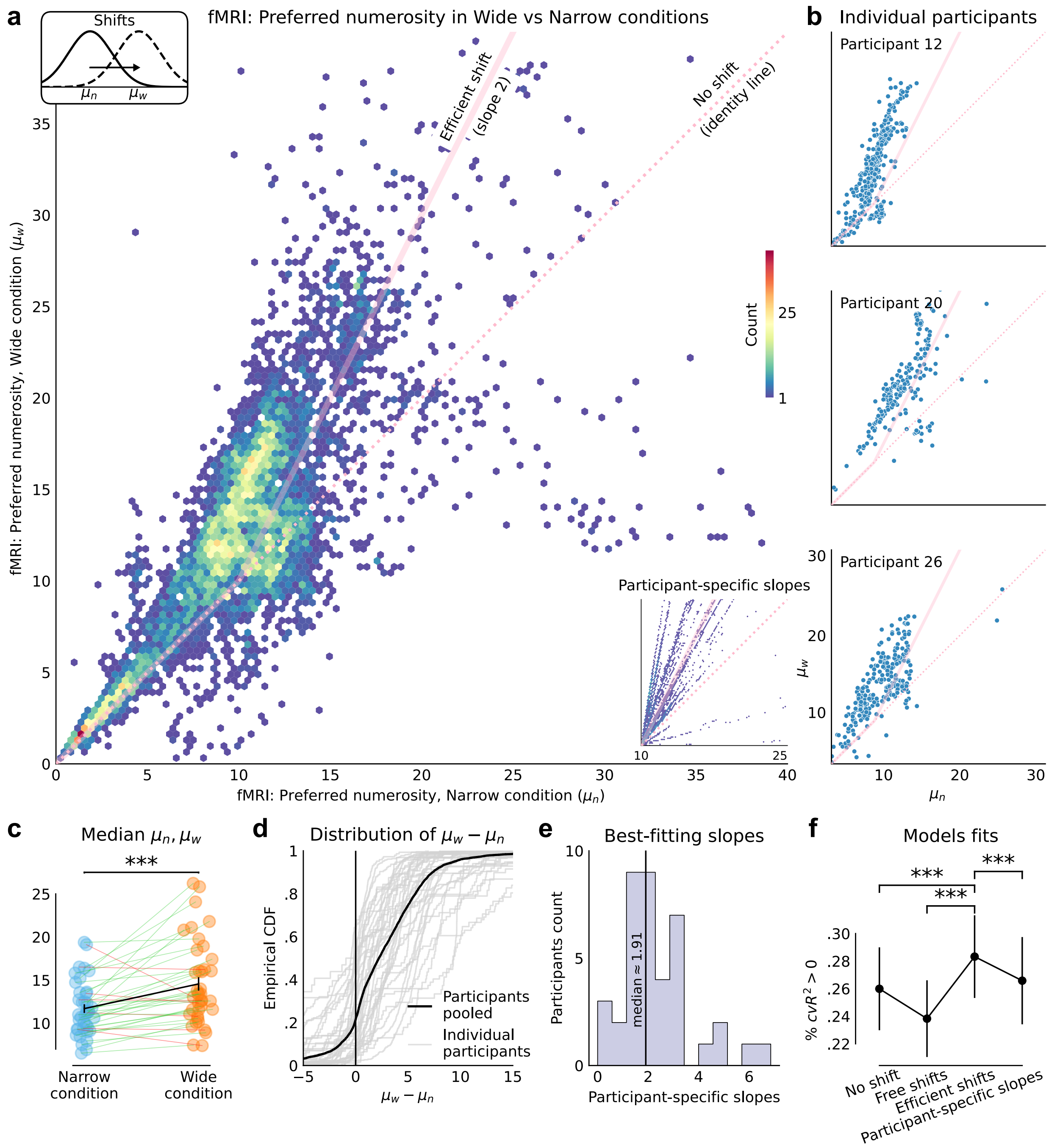


**Fig. S6: Ground-truth fit: Efficient shifts of neural receptive fields across priors.** As Figure 3 in the main text, but with receptive-field models fit using the ground-truth numbers rather than the responses.

$P = 0.0014$.

- For 32 out of 39 participants, the across-voxels median preferred numerosity is larger in the Wide condition than in the Narrow condition (Fig. S6c; across-participants paired t-test of equality of the medians in the two conditions: $t(38) = 4.31$, $P = 1.1 \times 10^{-4}$).

Overall, the preferred numerosities of 78% of voxels increase in the Wide condition as compared to the Narrow condition (Fig. S6d).

- $\mu_w$ for voxels with $\mu_n \in [14.5, 15.5]$ is on average $20.45$; sem: 0.50.
- 2.4% of voxels have preferred numerosities greater than $25$ in the Narrow condition.
- 'Efficient-shift' model compared to 'free-shift' model: across-participants paired t-test of equality of the proportions of voxels with cv$R^2 > 0$: $t(38) = 8.34$, $P = 4 \times 10^{-10}$. Compared to 'no-shift' model: $t(38) = 4.21$, $P = 1.5 \times 10^{-4}$. Compared to 'participant-specific slopes' model (lower-right inset in Fig. S6a): $t(38) = 3.86$, $P = 4.3 \times 10^{-4}$. See Figure S6f.
- Slope parameter in the 'participant-specific slopes' model: across-participant median: $1.91$, mean: $2.32$, sem: $0.24$ (Fig. S6e). The mean best-fitting value is significantly different from 1 (t-test $t(38) = 5.59$, $P = 2 \times 10^{-6}$), but not from 2 ($t(38) = 1.36$, $P = 0.18$).

**Receptive fields widths**

- Correlation between the widths parameters $\sigma_n$ and $\sigma_w$ in the two conditions: participants pooled: $r = 0.27$, $P = 8 \times 10^{-131}$, $N = 7977$; across participant: average $r = 0.45$, IQR: $0.25$-$0.60$.
- For 31 out of 39 participants, the median width (across voxels) in the Wide condition is larger than in the Narrow condition (paired t-test of equality: $t(38) = 4.9$, $P = 2 \times 10^{-5}$; Fig. S7c). Overall, the widths of 81% of voxels increase in the Wide condition as compared to the Narrow condition (Fig. S7d).
- Scale parameter of the 'participant-specific scaling' model: across-participants median: 1.3, median: 1.3, sd: 0.25 (Fig. S7e). The mean best-fitting value is significantly greater than 1 (t-test $t(38) = 7.23$, $P = 1.2 \times 10^{-8}$), and significantly lower than 2 ($t(38) = 17.9$, $P = 4 \times 10^{-20}$.
- 'Fixed scaling' model ($r_\sigma = 1.3$) compared to 'participant-specific scaling' model: across-participants paired t-test of equality of the proportions of voxels with cv$R^2 > 0$: $t(38) = 3.5$, $P = 1.3 \times 10^{-3}$. Compared to 'free widths' model: $t(38) = 9.0$, $P = 7 \times 10^{-11}$. Compared to 'no width change' model: $t(38) = 3.4$, $P = 1.6 \times 10^{-3}$. See Figure S7f.

In short, the results obtained with the receptive fields models fit with the correct numbers are qualitatively identical, and quantitatively very similar, to those obtained when fitting with the participants' estimates.

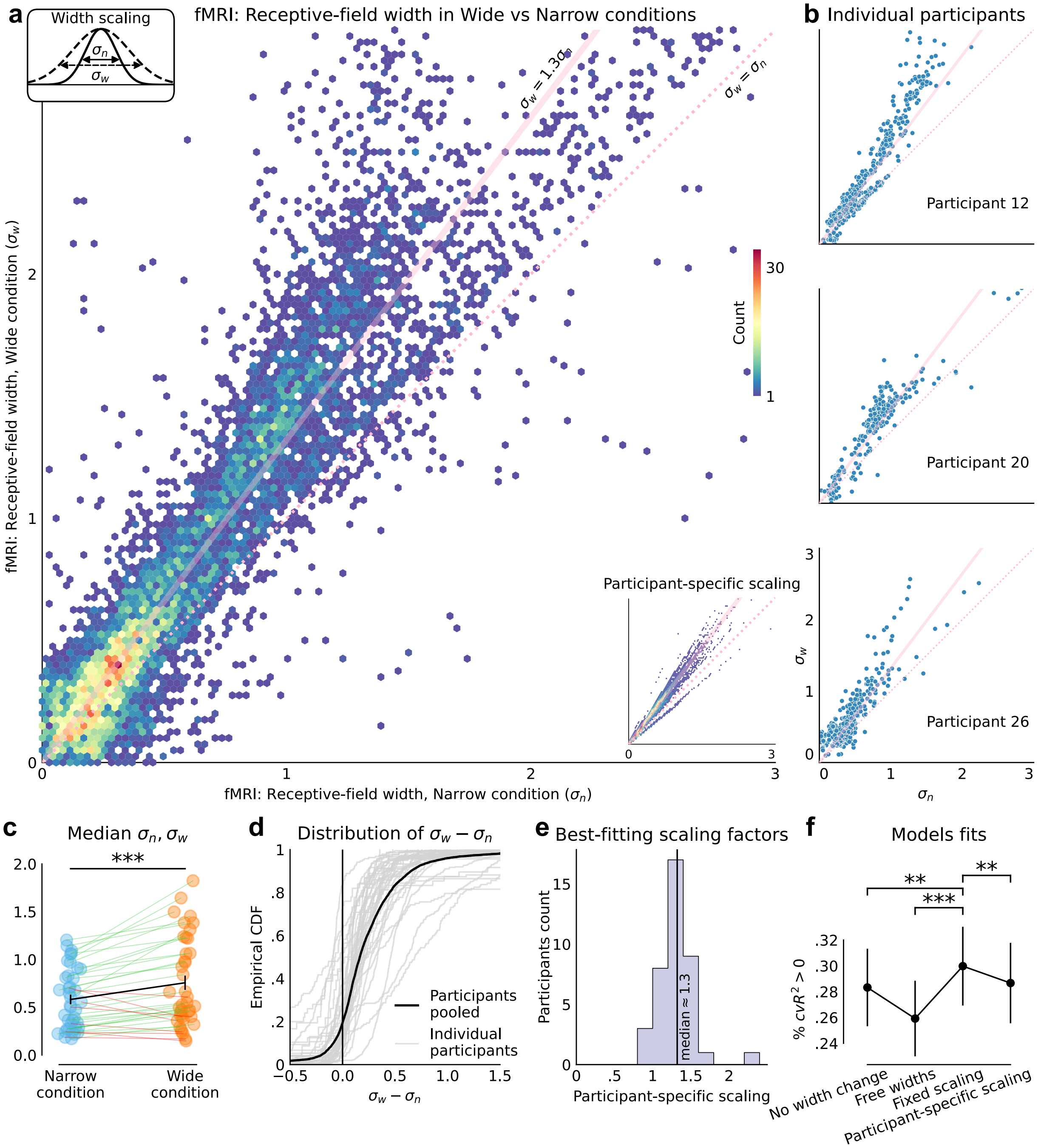


**Fig. S7: Ground-truth fit: The receptive fields broaden under the wider range.** As Figure 4 in the main text, but with receptive-field models fit using the ground-truth numbers rather than the responses.

### 4 nPRF estimation checks

#### Parameter recovery confirms unbiased nPRF estimation across stimulus ranges

A potential concern with comparing nPRF parameters across conditions with different stimulus ranges is that the model fitting procedure itself might be biased by the range of numerosities presented. To address this, we performed model recovery analyses for both condition-linking parameters used in the main analyses: the shift in preferred numerosity ($r_\mu$) and the scaling of tuning width ($r_\sigma$) (see Methods in main text for details). Briefly, for each parameter we simulated voxels under a $2 \times 2$ factorial design, crossing two generating values ($r_\mu = 1$, no shift, or $r_\mu = 2$, full range adaptation; and, for the width analysis, $r_\sigma = 1$, no widening, or $r_\sigma = 1.29$, the empirical group-average widening) with two stimulus designs (full, matching our actual experiment; censored, restricted to the shared range $[10, 25]$). Generating tuning parameters (preferred numerosity and width) were not chosen arbitrarily but resampled, as pairs, from the empirical encoding-model fits in right parietal cortex (8,785 voxels from 39 participants), and simulated responses were corrupted with Gaussian noise calibrated so that the median single-trial $R^2$ of the simulated voxels matched the empirical median. We repeated each condition 100 times and applied the same nPRF fitting procedure used in the main analyses. As an additional robustness check, we repeated the entire analysis with a subject-wise sampling scheme, in which each iteration simulated all supra-threshold voxels of one randomly drawn participant (29–535 voxels, mirroring the per-participant structure of the real analysis) rather than a fixed pool of 250 voxels drawn across participants; this is shown as a second row in Fig. S8 and Fig. S9.

**Shift parameter** As shown in Fig. S8, the recovered shift parameter $r_\mu$ closely matched the generating value in all four conditions, under the pooled sampling scheme (recovered $r_\mu = 1.00 \pm 0.04$ for the null and $2.00 \pm 0.05$ for the full-shift condition, mean $\pm$ SD across 200 simulated datasets per generating value, pooling both designs). Critically, restricting the stimulus range to the shared interval (censored design) did not introduce any systematic bias relative to the full design, confirming that differences in stimulus sampling between the Narrow and Wide conditions cannot explain the tuning shifts reported in the main text. Under the subject-wise sampling scheme, recovery was somewhat noisier, as expected given the smaller and more variable per-iteration voxel counts, and showed a small downward bias for the full-shift condition ($r_\mu = 1.03 \pm 0.16$ and $1.93 \pm 0.24$ for the null and full-shift conditions, respectively); the direction and magnitude of the shift were nonetheless correctly recovered in every condition. Code for this simulation is available at `https://github.com/ruffgroup/neural_priors/blob/main/neural_priors/revision%20feb%202026/simulate_data.py`.

**Width-scaling parameter** Similarly, when we simulated and re-estimated the factor scaling tuning width between the narrow and wide conditions ($r_\sigma$; Fig. S9), we found very consistent recovery: under the pooled sampling scheme, the recovered scaling factor was $r_\sigma = 0.99 \pm 0.03$ for the null condition and $1.26 \pm 0.02$ for the empirically observed widening

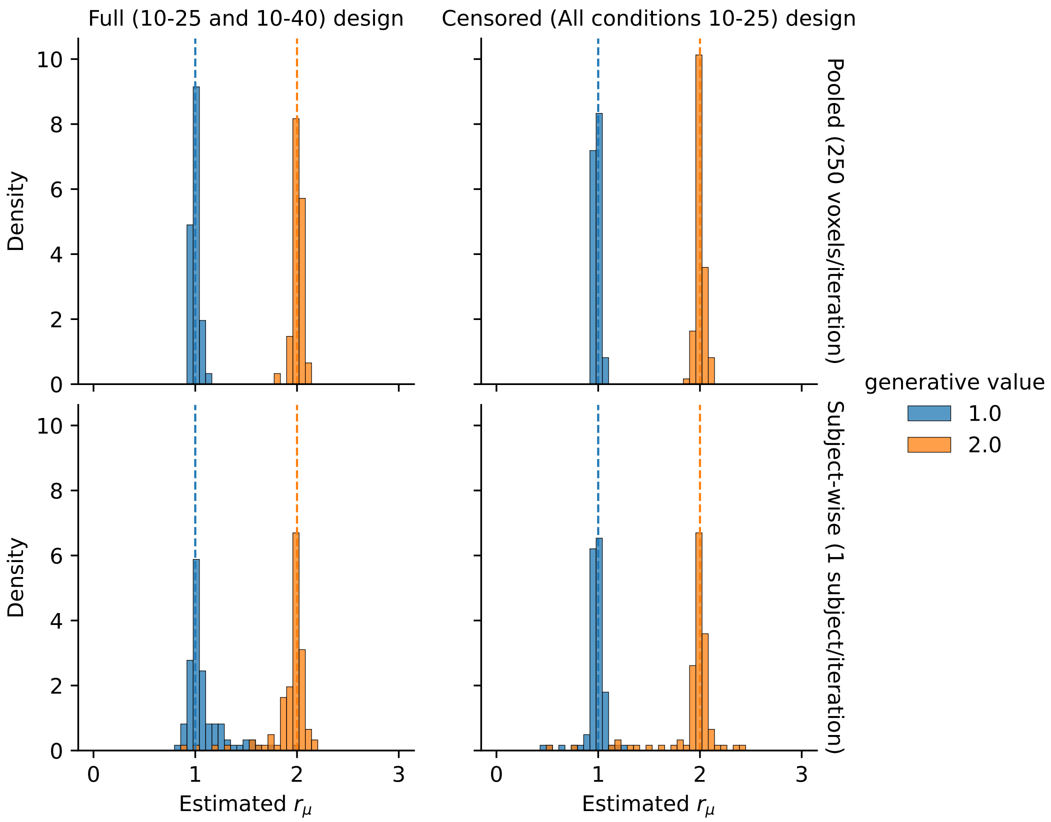


**Fig. S8: Parameter recovery analysis confirms unbiased recovery of the shift in preferred numerosity across stimulus ranges.** Distributions of the recovered shift parameter $r_\mu$ (shared across voxels, freely fitted), over 100 simulated datasets per condition for each of two sampling schemes: a fixed pool of 250 voxels pooled across participants (top row), and, as a robustness check, all supra-threshold voxels of one randomly drawn participant per iteration (bottom row). Columns: full design (narrow range 10–25, wide range 10–40, left) and censored design (both ranges 10–25, right). Blue: generative $r_\mu = 1$ (no shift); orange: generative $r_\mu = 2$ (full shift); dashed vertical lines mark the generative value. Pooled sampling: recovered $1.00 \pm 0.04$ and $2.00 \pm 0.05$ (mean $\pm$ SD, pooling both designs). Subject-wise sampling: recovered $1.03 \pm 0.16$ and $1.93 \pm 0.24$. In every panel the recovered distributions are centred close to the generative value and the two generative conditions are well separated, confirming that nPRF model fitting is not systematically distorted by differences in stimulus range or sampling scheme between conditions.

($r_\sigma = 1.29$); recovery was again somewhat noisier, but still centred on the generating value, under the subject-wise sampling scheme ($0.99 \pm 0.05$ and $1.27 \pm 0.06$, respectively). As for the shift analysis, restricting the stimulus range to the shared interval did not introduce systematic bias. Together, these recovery analyses confirm that neither the shift in preferred numerosity nor the change in tuning width reported in the main text can be explained by the fitting procedure itself or by differences in stimulus sampling between conditions. Code for this simulation is available at `https://github.com/ruffgroup/neural_priors/blob/main/neural_priors/revision%20aug%202026/simulate_data_sd.py`.

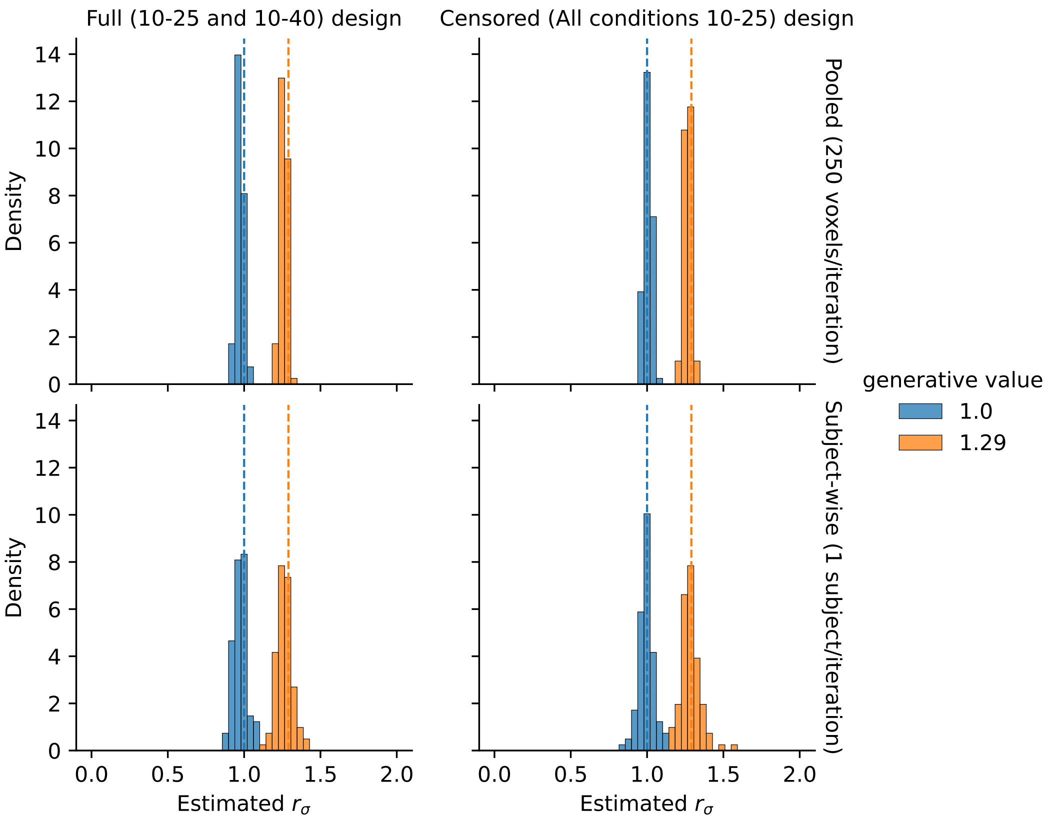


**Fig. S9: Parameter recovery analysis confirms unbiased recovery of the change in tuning width across stimulus ranges.** As Fig. S8, but for the width-scaling factor $r_\sigma$. Blue: generative scaling factor 1 (no widening); orange: generative scaling factor 1.29, the empirical group-average value; dashed vertical lines mark the generative value. Pooled sampling: recovered $0.99 \pm 0.03$ and $1.26 \pm 0.02$ (mean $\pm$ SD, pooling both designs). Subject-wise sampling: recovered $0.99 \pm 0.05$ and $1.27 \pm 0.06$. The recovered distributions are centered on the generative values under both sampling schemes — with, if anything, a slight underestimation of the true widening in the pooled scheme — and do not overlap. Thus, the widening of tuning curves in the wide condition reported in the main text is neither produced nor inflated by the fitting procedure, despite the presence of many voxels with preferred numerosities below the stimulus range, for which individual tuning widths are only weakly constrained.

**Widths** The results of the recovery procedure for the width parameter, which may be difficult to estimate reliably, are shown in Figure S10. The Spearman correlation between generative and recovered per-voxel tuning widths is $\rho = 0.60$ ($n = 194,212$). This correlation is higher for voxels whose preferred numerosity falls within the presented stimulus range ($\rho = 0.70$, $n = 104,640$) than for those outside it ($\rho = 0.50$, $n = 89,572$), where the response profile is not fully sampled. But even for these out-of-range voxels, the correlation remains considerably above zero, indicating that individual tuning-width estimates retain real information even when the response profile is monotonic within the stimulus set.

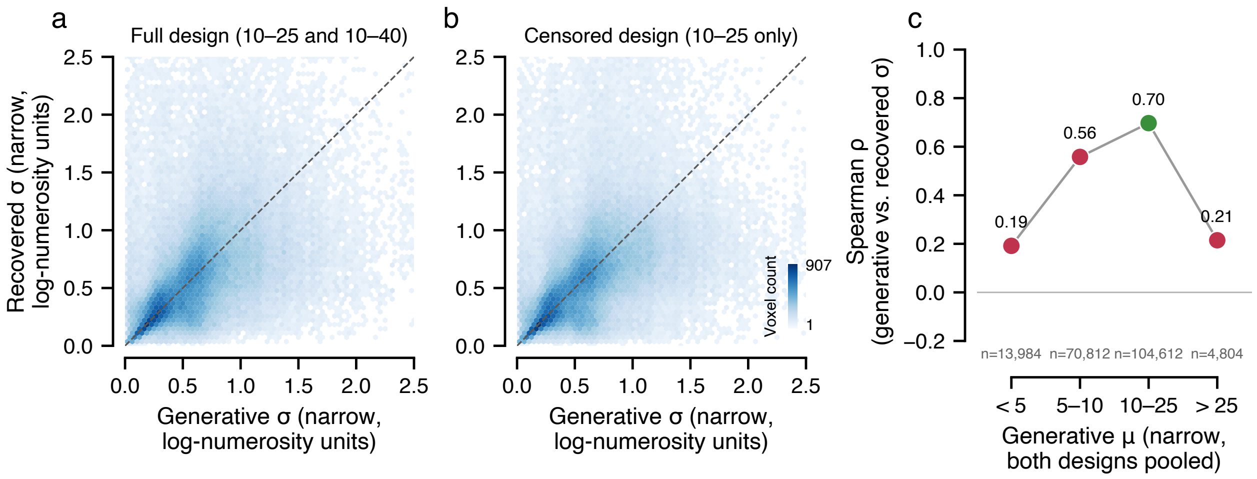


**Fig. S10: Width-parameter recovery. a,b,** Recovered width parameter, $\sigma$, as a function of the generative parameter, with the full design (a) and the censored design (b). **c,** Spearman correlation between the recovered and generative parameter, as a function of the preferred-numerosity parameter, $\mu$.

**Preferred numerosities** Although the estimation of the preferred-numerosity parameter is generally more reliable than that of the width parameter, a difficulty arises when the preferred numerosity is below 10, i.e., outside of the range of numerosities that are actually presented in the experiment. Looking at the preferred-numerosity estimates that are below 10, we note that the Pearson correlation between these estimates across conditions is 0.81, and that a Deming regression (which assumes noise in both the dependent and the independent variables) yields a slope of 1.098 and an intercept of -0.011, two values close to the identity function (Fig. S11a). These preferred-numerosity estimates are thus consistent across conditions. This argues against the hypothesis that estimates below 10 are completely arbitrary or dominated by fitting noise, but it does not establish that they are unbiased estimates of the true preferred numerosities (as the same estimation bias could occur in both conditions). It suggests, however, that the encoding properties of these populations are stable, across conditions.

The recovery simulations detailed above, which include voxels with $\mu < 10$ in their empirical proportions, allow us to more closely examine the quality of estimates. For voxels whose generative preferred numerosity lay within the presented range (10–25), preferred numerosity was recovered accurately (rank correlation between generative and recovered values $\rho = 0.79$; median absolute error 0.74; median bias 0.05). For voxels with generative preferred numerosities just below the range (5–10), recovery was less precise ($\rho = 0.20$), with a small positive bias (median +0.38 across all out-of-range voxels, reflecting regression toward the range boundary), although the median absolute error was only 1.96. The fits correctly identified 71% of below-range voxels as preferring a numerosity below 10. For generative preferred numerosities far below the range, estimates were strongly biased towards the presented range and carried little information about the true value (Fig. S11b,c).

In other words, the fitting procedure reliably recognizes that a voxel prefers a numerosity

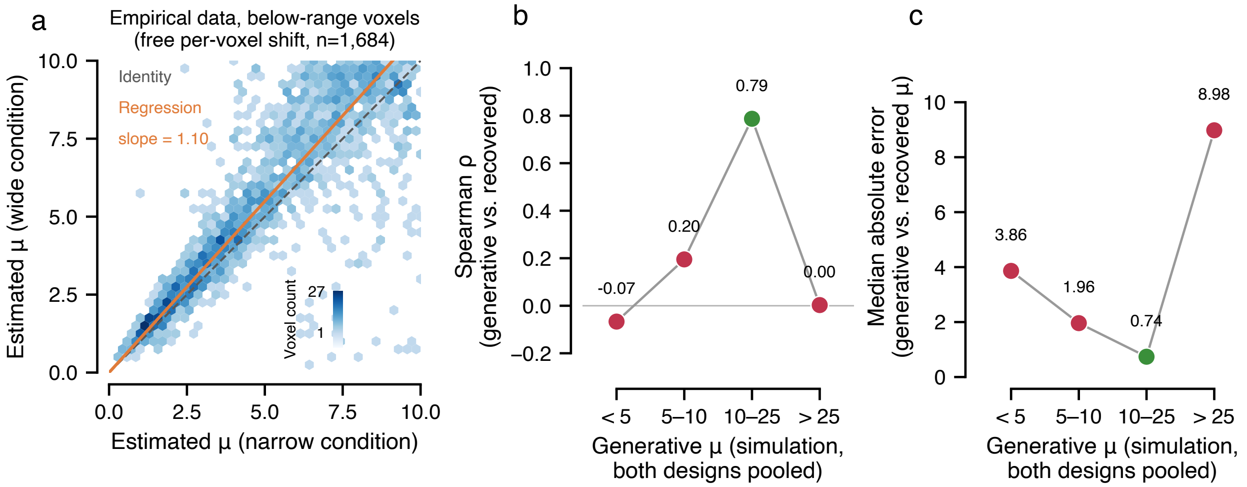


**Fig. S11: Preferred-numerosity parameter analysis. a,** Estimated preferred numerosities in the wide vs. narrow condition, $\mu_w$ and $\mu_n$, when these are below 10, with Deming regression and identity line. **b,c,** Spearman correlation (b) and median absolute error (c) between the recovered and generative parameter $\mu$, binned per values of the generative parameter.

below the presented range, but not precisely where below the range its preference lies. Overall, the apparent stability below 10 is compatible with (rather than direct evidence for) our conjecture that these receptive fields remain unchanged across conditions. This does not affect our main conclusions: the efficient-coding model makes predictions about the rescaling of tuning for preferred numerosities within the presented ranges, and the recovery of the population-level rescaling parameters (shift and width scaling; Supplementary Figs. S8 and S9) is accurate despite the presence of these below-range voxels.

#### Preferred numerosities and widths with censored fits

One might argue that the shift in preferred numerosities might (partly) arise due to the fact that in the Wide condition, the encoding model is fit to a broader range of stimuli, potentially biasing preferred numerosities toward more extreme values. To rule out this possibility, we re-fit the 'Free shifts' model and the 'Participant-specific shifts' model, restricting the input numerosities to the range shared between the Narrow and Wide conditions (henceforth *censored* fits). Note that these censored fits are necessarily noisier, as the Wide condition then contains only half the amount of data. Reassuringly, the shift in median preferred numerosity remains highly significant under the censored fits (paired $t$-test: $t(38) = 5.60$, $P = 2 \times 10^{-6}$), and the slope parameter remains distributed (across participants) around 2 (median: 1.71, mean: 2.04, sem: 0.23), with a mean significantly different from 1 ($t(38) = 4.55$, $P = 5 \times 10^{-5}$), but not from 2 ($t(38) = 0.18$, $P = 0.86$), closely mirroring the results from the uncensored fits. Figure S12 shows the obtained preferred numerosities, which exhibit the same patterns as with the uncensored fits. These results confirm that the reported tuning shifts are not an artifact of differential stimulus sampling between conditions.

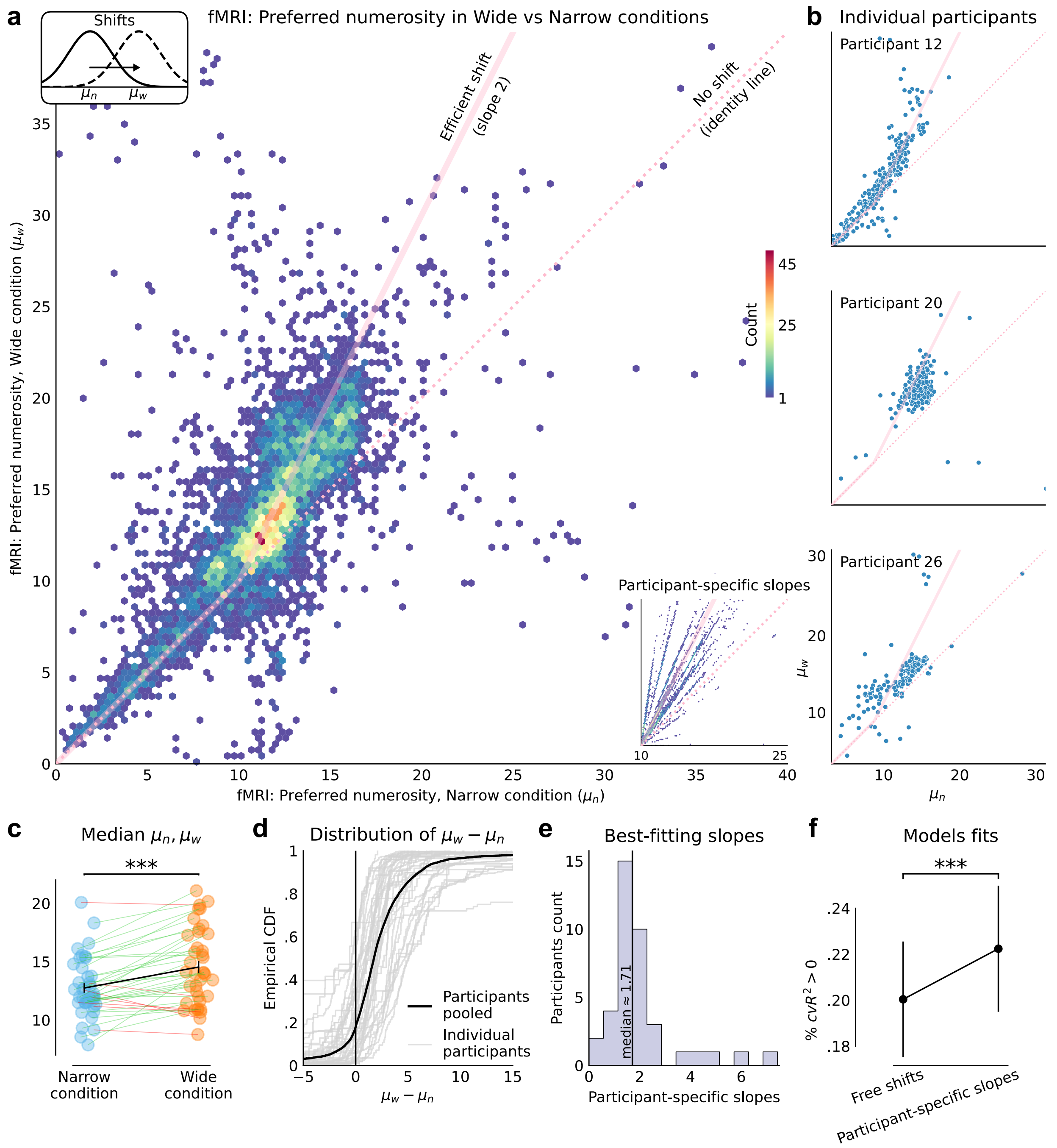


**Fig. S12: Censored fits: preferred numerosities.** As Figure 3 in the main text, but with fits restricted to the range shared between the Narrow and Wide conditions.

We conduct the same censored-fits analysis for the width parameter. Specifically, we fit the 'Efficient shifts, free widths' model and the 'Efficient shifts, participant-specific width-scaling' model on data restricted to the range shared between the Narrow and Wide conditions. Figure S13 shows the obtained widths, which exhibit the same patterns as with the uncensored fits. With the free-widths model, the two widths are significantly correlated (participants pooled: Spearman's $\rho = 0.80$, $P < 10^{-320}$, $N = 6586$; across participant: average $\rho = 0.66$, IQR: $0.59$-$0.85$). For a majority of participants (30 out of 39), the median width (across voxels) in the Wide condition is larger than in the Narrow condition (paired t-test of equality: $t(38) = 3.74$, $P = 6 \times 10^{-4}$; Fig. S13c). Overall, the widths of 68% of voxels increase in the Wide condition as compared to the Narrow condition (Fig. S13d).

With the model with participant-specific width-scaling, the scaling parameter $r_\sigma$ is significantly greater than 1 (t-test $t(38) = 4.16$, $P = 1.8 \times 10^{-4}$), and significantly lower than 2 ($t(38) = 8.6$, $P = 1.8 \times 10^{-10}$). Its across-participant average is 1.3 (median: 1.2, sem: 0.08; Fig. S13e). These results, consistent with those obtained with the uncensored fits, attest to the robustness of our parameter estimation, and substantiate our conclusions.

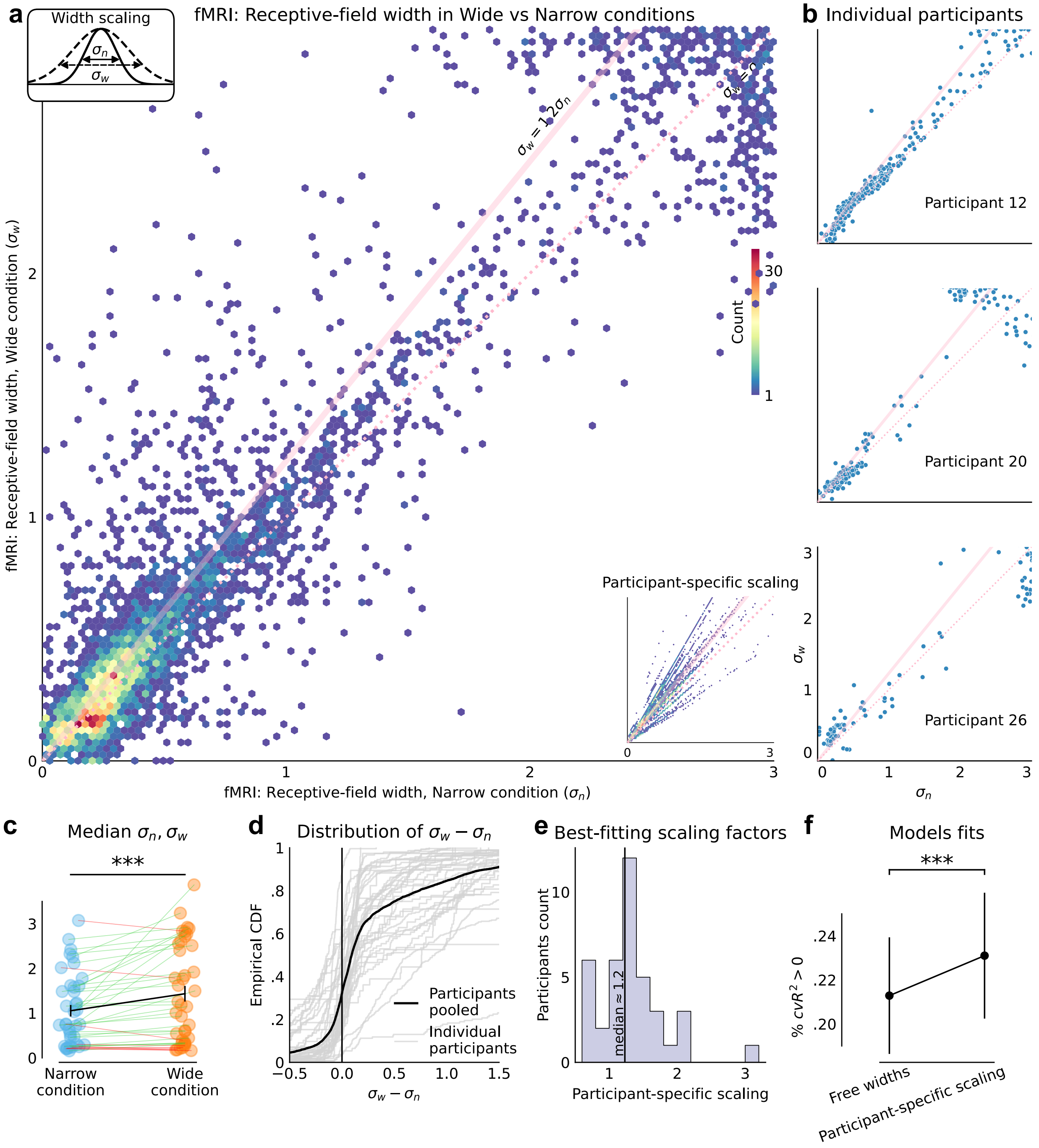


**Fig. S13: Censored fits: widths.** As Figure 4 in the main text, but with fits restricted to the range shared between the Narrow and Wide conditions.

### 5 Participants' responses and bias

Figure S14 shows the average responses provided by the participant in response to each numerosity, as well as their bias. The behavior of participants exhibit all the typical patterns found in magnitude estimation tasks (including with numerosity)[1–4]: overestimation of small magnitudes, underestimation of large magnitudes, a 'central tendency of judgment' wherein the crossover point (where the bias vanishes) depends on the width of the prior, and a negative bias for large numerosities that is larger than the positive bias for small numerosities. We also show the biases and standard deviations of individual participants in Figures S15 and S16.

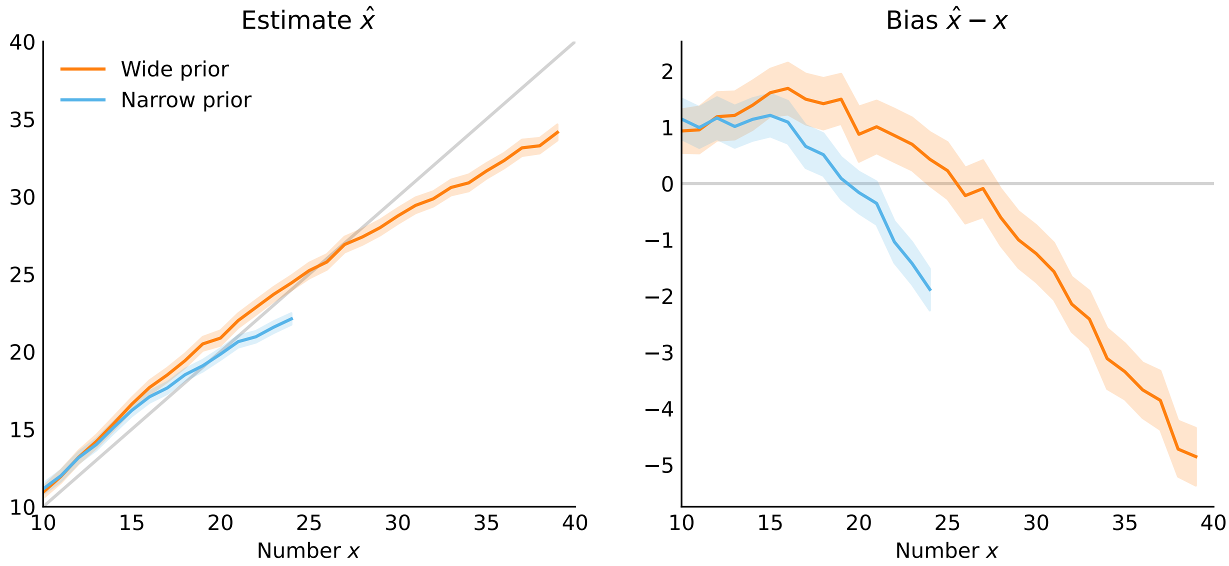


**Fig. S14: Participants' responses and bias.** Mean participants' response $\hat{x}$ (*left*), and bias $\hat{x} - x$ (*right*), as a function of the presented number $x$. Shaded areas show the 5%-95% credible intervals (see Methods for details on the statistical estimation procedure).

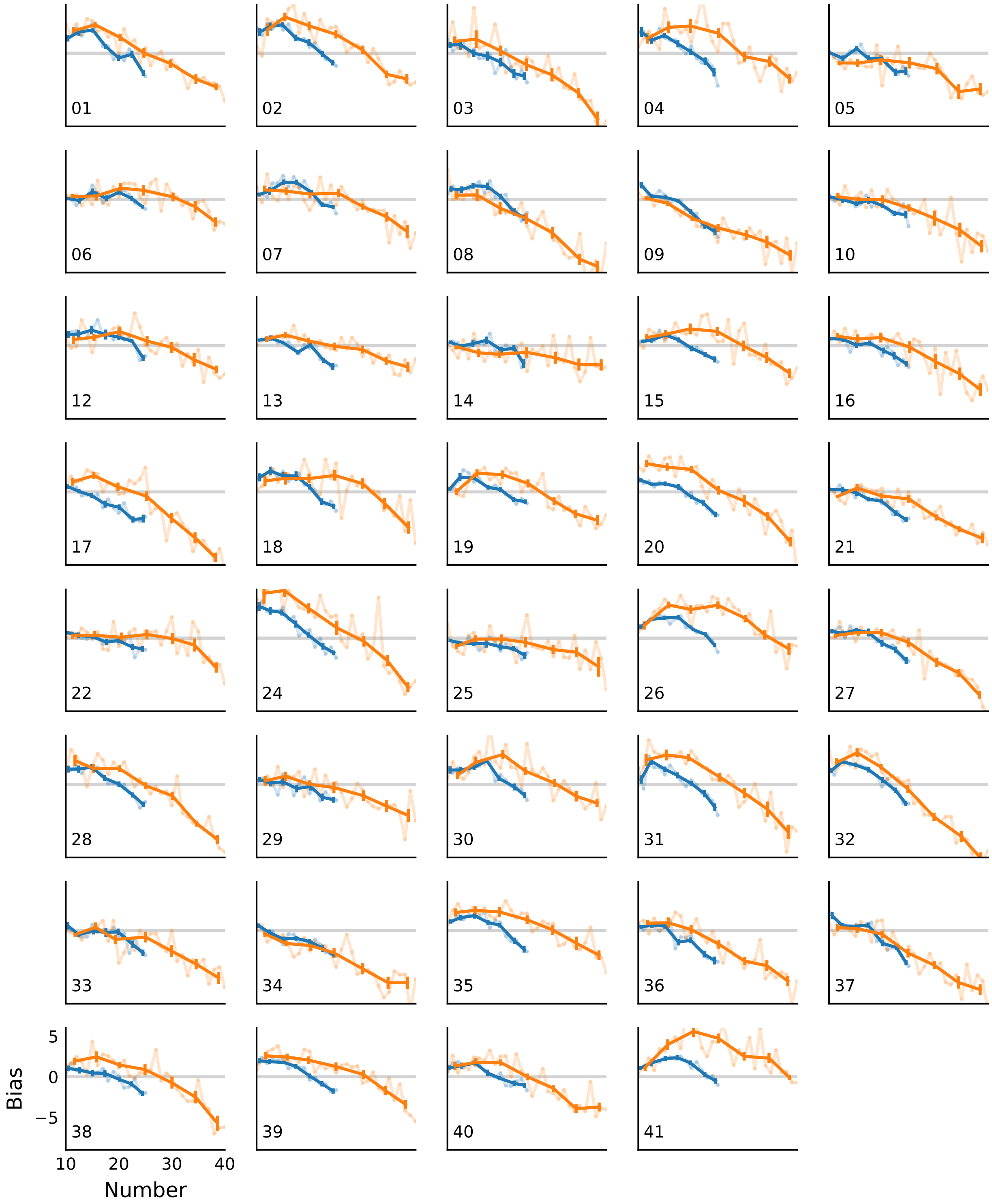


**Fig. S15: Responses of individual participants.** Empirical average of each participant's bias (difference between the response and the presented number) as a function of the presented number; with numbers pooled in bins (thick lines) or not (thin lines). Error bars: sem.

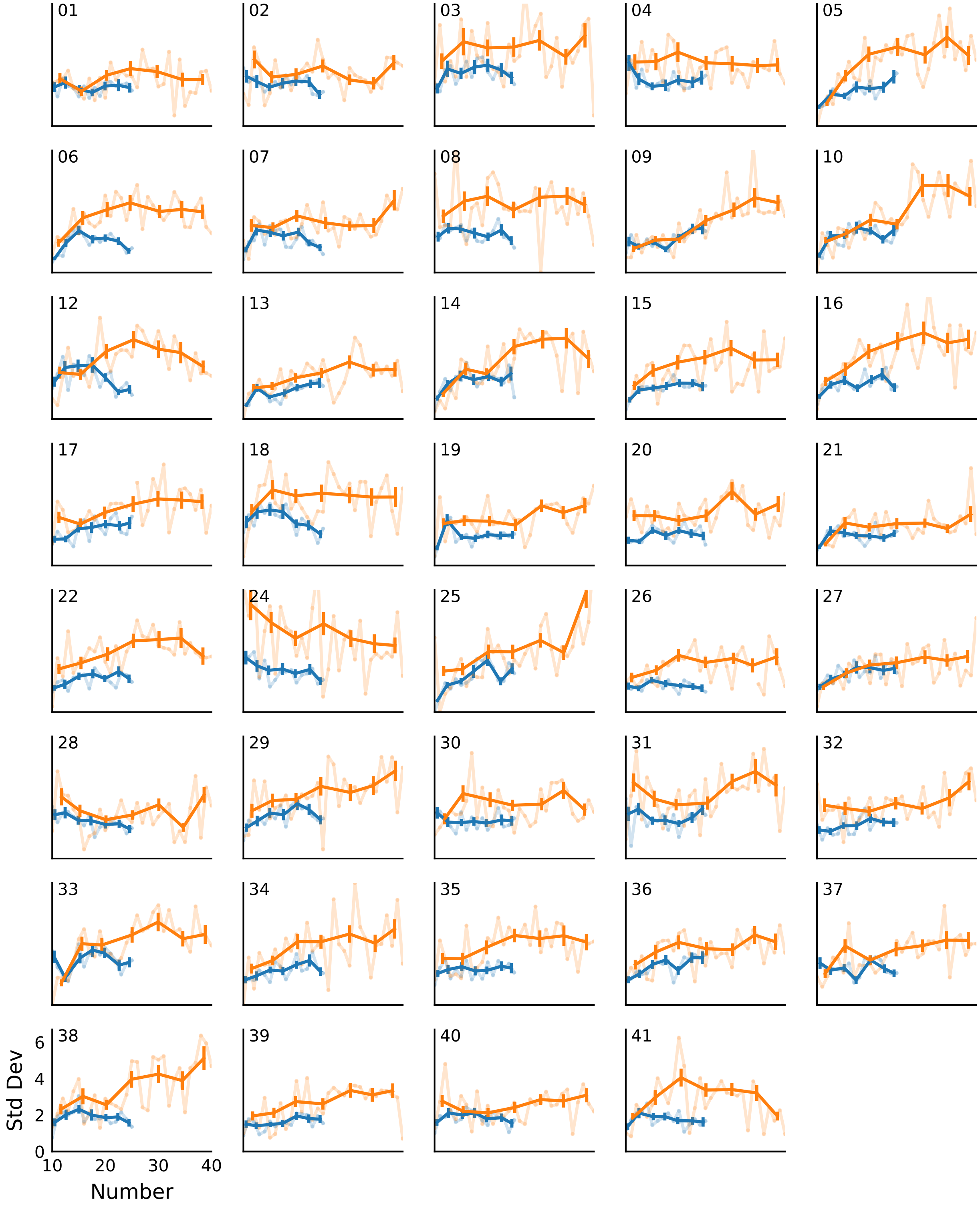


**Fig. S16: Standard deviations of individual participants.** Empirical standard deviation of each participant's responses as a function of the presented number; with numbers pooled in bins (thick lines) or not (thin lines). Error bars: sem.

### 6 Retinotopic account fails to account for nPRF shifts

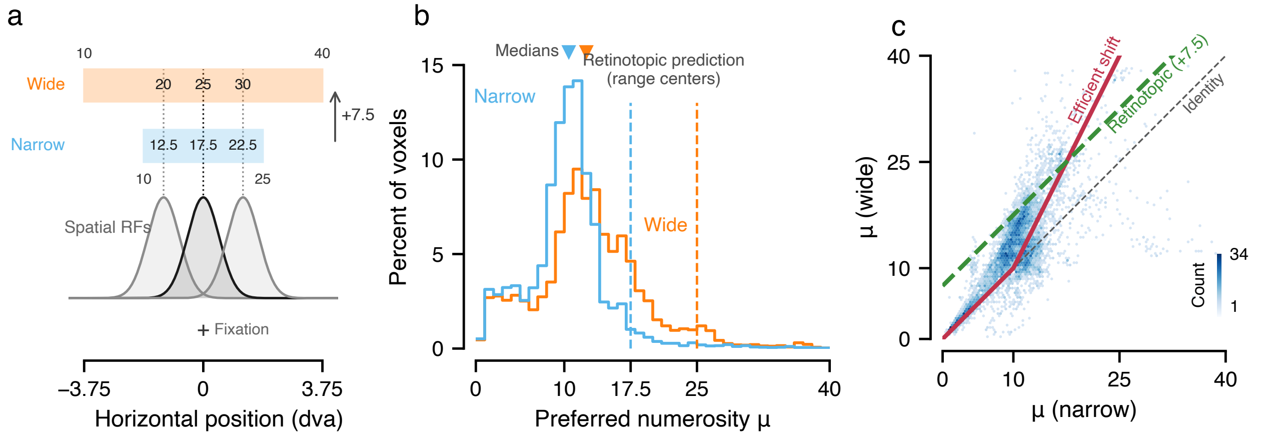


**Fig. S17: Numerosity tuning does not reflect the retinotopic position of the response bar.** **a**, The retinotopic (bar-position) account. Both response bars were centered on fixation and used the same spatial scale (0.25 degrees of visual angle per numerosity unit), so the narrow bar (10–25) spanned 3.75 and the wide bar (10–40) spanned 7.5 degrees of visual angle. A voxel with a spatial receptive field at a fixed horizontal position (Gaussian profiles; dotted lines) therefore reads out a numerosity that is exactly 7.5 units higher on the wide than on the narrow bar, for every position — e.g. a foveal receptive field (black) maps to 17.5 (narrow) and 25 (wide), the centers of the two ranges. Bar-position tuning thus predicts (i) preferred numerosities concentrated at the range centers, given the foveal bias of parietal spatial receptive fields, and (ii) a constant *additive* shift of $+7.5$ between conditions. **b**, Distributions of preferred numerosities ($\mu$) across voxels in the right parietal region of interest for the narrow (blue) and wide (orange) conditions, estimated with the model variant in which the shift in preferred numerosity is fitted freely for every voxel, so that $\mu$ is effectively unconstrained in both conditions (7,264 voxels with positive cross-validated $R^2$; 39 participants). Both distributions peak near the shared lower range bound, with medians of 10.5 and 12.5 (triangles), far below the retinotopically predicted central tendencies (dashed lines at 17.5 and 25); only $\sim$6% of voxels prefer a numerosity above the respective range center. **c**, Preferred numerosity in the wide against the narrow condition for the same voxels (hexagonal bins; color denotes voxel count). Green dashed line: the additive retinotopic prediction $\mu_w = \mu_n + 7.5$ from panel a. Red line: the efficient-coding (range adaptation) prediction of a *multiplicative* remapping anchored at the shared lower bound, $\mu_w = 2\,(\mu_n - 10) + 10$ for $\mu_n \geq 10$ and no shift below the presented range (thin gray dashed line, identity). The data follow the multiplicative pattern: the efficient-coding prediction yields a smaller absolute prediction error than the retinotopic prediction in 66.6% of voxels (median absolute error 2.8 vs. 5.5) and in 31 of 39 participants (Wilcoxon signed-rank test on per-participant median absolute errors, $W = 75$, $P = 1.7 \times 10^{-6}$). Consistent with a multiplicative rather than additive shift, the per-voxel shift $\mu_w - \mu_n$ increases with $\mu_n$ within the presented range (per-participant regression slope, mean 0.38; $t(38) = 2.91$, $P = .006$).
